# Reappraisal of Spinach-based RNA Visualization in Plants

**DOI:** 10.1101/2020.09.24.310607

**Authors:** Zhiming Yu, Fengling Mei, Haiting Yan, Qiyuan Chen, Mengqin Yao, Shuyue Liu, Yue Wang, Xian Zhang, Pengcheng Zhang, Stephen Jackson, Nongnong Shi, Yule Liu, Yiguo Hong

**Author notes:** The author responsible for distribution of materials integral to the findings presented in this article in accordance with the policy described in the instructions for Authors (www.plantphysioll.org) is: Yiguo Hong (, and). **Author Contributions:** Z.Y. designed, performed all experiments, analyzed data and drafted the manuscript; F.M. H.Y. and Q.C. performed plant transformation, plant RNA extraction and RT-PCR analysis. M.Y., S.L., Y.W., X.Z., and P.Z. performed virus-based RMG assays; S.J., N.S., and Y.L. were involved in the analysis of data and helped write the article; Y.H. initiated the project, conceived experiments, analyzed data, and wrote the article. **Competing interests:** The authors declare no competing interests.

## Abstract

RNAs can be imaged in living cells using molecular beacons, RNA-binding labeled proteins and RNA aptamer-based approaches. However, Spinach RNA-mimicking GFP (RMG) has not been successfully used to monitor cellular RNAs in plants. In this study, we re-evaluated Spinach-based RNA visualization in different plants via transient, transgenic, and virus-based expression strategies. We found that like bacterial, yeast and human cellular tRNAs, plant tRNAs such as tRNA^Lys^ (K) can protect and/or stabilize the spinach RNA aptamer interaction with the fluorophore DFHBI enabling detectable levels of green fluorescence to be emitted. The tRNA^Lys^-spinach-tRNA^Lys^ (KSK), once delivered into “chloroplast-free” onion epidermal cells can emit strong green fluorescence in the presence of DFHBI. Transgenic or virus-based expression of monomer KSK, in either stably transformed or virus-infected *Nicotinana benthamiana* plants, failed to show RMG fluorescence. However, incorporating tandem repeats of KSK into recombinant viral RNAs, enabled qualitative and quantitative detection, both in vitro and ex vivo (ex planta), of KSK-specific green fluorescence, though RMG was less obvious in vivo (in planta). These findings demonstrate Spinach-based RNA visualization has the potential for *ex vivo* and *in vivo* monitoring RNAs in plant cells.

**One sentence summary:** Spinach-based RMG technology was reevaluated to have potential for ex vivo and in vivo monitoring RNAs in plant cells.

## INTRODUCTION

RNAs primarily act as messengers to convey genetic information from DNA to protein. However, the functionalities of RNAs are much broader. Increased evidence has demonstrated that RNAs can be potent regulators modulating gene expression at transcriptional, post-transcriptional and translational levels (Shi et al., 2008; Zhang et al., 2019). In plants, cellular mRNAs, small interfering RNA (siRNA), microRNAs, and pathogenic viral and viroid RNAs, can move from cell to cell through plasmodesmata and spread to distal tissues via the phloem superhighway (Carrington et al., 1996; Xoconostle-Cázares et al., 1999; Yoo et al., 2004; Buhtz et al., 2008; Deeken et al., 2008; Uddin and Kim, 2013; Chen et al., 2018; Shahid et al., 2018). Some of these mobile RNAs function as intra- and intercellular as well as systemic signals to control plant defense, growth and development (Jackson and Hong, 2012; Liu and Chen, 2018; Zhang et al., 2019). For instance, mobile *Gibberellic acid insensitive* mRNA regulates leaf morphology in *Arabidopsis*, tomato and pumpkin (Haywood et al., 2005). Systemic trafficking of *CmNACP* affects shoot and root apical meristem maintenance in pumpkin (Ruiz-Medrano et al., 1999). *BEL5* mRNA moves from leaf to stolon tip to promote tuber formation and development in potato (Banerjee et al., 2006). Transportable *AtFT* (Li et al., 2009; Li et al., 2011; Lu et al., 2012; Luo et al., 2018; Ellison et al., 2020), *FVE* (Yang and Yu, 2010), *AGL24* (Yang and Yu, 2010) and *ATC* (Huang et al., 2012) mRNAs regulate flowering in *Arabidopsis*. Furthermore, many RNAs are also able to move across hetero-graft scions between different plants (Notaguchi et al., 2015) and ecotypes (Thieme et al., 2015), between parasitic plant and its hosts in a bidirectional manner, or even between plants and fungi (Kim et al., 2014; Uddin and Kim, 2013). Thus, mobile RNAs have enormous potentials in regulating plant growth and development and in response to biotic or abiotic stresses and (Jackson and Hong, 2012; Thieme et al., 2015; Liu and Chen 2018; Zhang et al., 2019). These emerging frontiers in RNA physiology demand the development of novel technologies to visualize RNAs in plants.

RNAs can be imaged in living cells using molecular beacons (MBs), RNA-binding labeled proteins (RBLPs) and RNA aptamer-based approaches (Tutucci et al., 2018). MBs involve a specific probe that perfectly complements with the target RNA in homogeneous solutions (Tyagi et al., 1996). RBLPs, such as MS2 (Bertrand et al., 1998), PUM-HD (Wang et al., 2002; Yamada et al., 2011; Filipovska et al., 2012; Yoshimura et al., 2012), hnRNPA1 (Scheiba et al., 2014), λN22 (Schönberger et al., 2012), Cas9 (Nelles et al., 2016) and Cas13a (Abudayyeh et al., 2017), bind to a specific sequence for detection. Unlike MB- or RBLP-based RNA assays, RNA aptamer Spinach (known as 24-2 or 24-2min), and its derivative Spinach2 mimic the Green Fluorescent Protein (GFP) when visualizing targeted RNAs (Paige et al., 2011; Strack et al., 2013; You and Jaffrey, 2015). These RNA aptamers bind to the fluorophore DFHBI (3,5-difluoro-4-hydroxybenzylidene imidazolinone) and form an intramolecular G-quadruplex to emit green fluorescence (Warner et al., 2014; Huang et al., 2014). This technology has been successfully used to directly monitor RNAs in bacterium (Paige et al., 2011; Pothoulakis et al., 2014; Zhang et al., 2015), yeast (Guet et al., 2015), and human cells (Paige et al., 2011); and to quantify cellular microRNAs (Huang et al., 2017). More recently, a similar fluorescent RNA aptamer dubbed Pepper has also been developed to image RNA in mammalian cells through its binding to the fluorophore ((4-((2-hydroxyethyl)(methyl)amino)-benzylidene)-cyanophenylacetonitrile) (Chen et al., 2019). However, use of fluorescent RNA aptamer-based RNA visualization has had little success in plants (Huang et al., 2012; 2017; Bai et al., 2020) although such techniques have attracted a great deal of attention from plant scientists (Ehrhardt and Frommer, 2012). In this study, we reevaluated the usefulness of ‘RNA-mimicking-GFP’ to monitor Spinach and Spinach-tagged RNAs via transient and transgenic expression, as well as virus-based delivery of recombinant RNAs, in different plant cells and tissues.

## RESULTS

### *In vitro* Spinach fluorescence

Prior to delivery of the Spinach RNA aptamer, or Spinach-tagged RNAs into plant cells and tissues, we tested if flanking a plant tRNA at both 5’ and 3’-ends of the Spinach RNA aptamer (Paige et al., 2011) would affect its binding to DFHBI and emission of green fluorescence (Fig. 1). We cloned the *Arabidopsis thaliana* lysine-tRNA (tRNA^Lys^) and Spinach (24-2min) (Paige et al., 2011) in the format of *AttRNA^Lys^-AttRNA^Lys^* (KK) or *AttRNA^Lys^-Spinach(24-2min)-AttRNA^Lys^* (KSK) into the pMD-19/T vector to generate pMD19-T/KK and pMD19-T/KSK constructs, respectively (Fig. 1A; Supplemental Data Set S1). The KK and KSK RNA transcription is driven by the T7 promoter in the two expression vectors. KK and KSK RNAs produced by *in vitro* transcription were readily detectable by agarose gel electrophoresis (Fig. 1B). Once mixed with DFHBI, only KSK RNA produced strong GFP-like green fluorescence (Fig. 1C-I). These data indicate that plant tRNA, similar to its bacterial, yeast or human counterpart (Paige et al., 2011; Pothoulakis et al., 2014; Guet et al., 2015; Zhang et al., 2015), can protect Spinach to enable GFP fluorescence *in vitro*.

**Figure 1.**
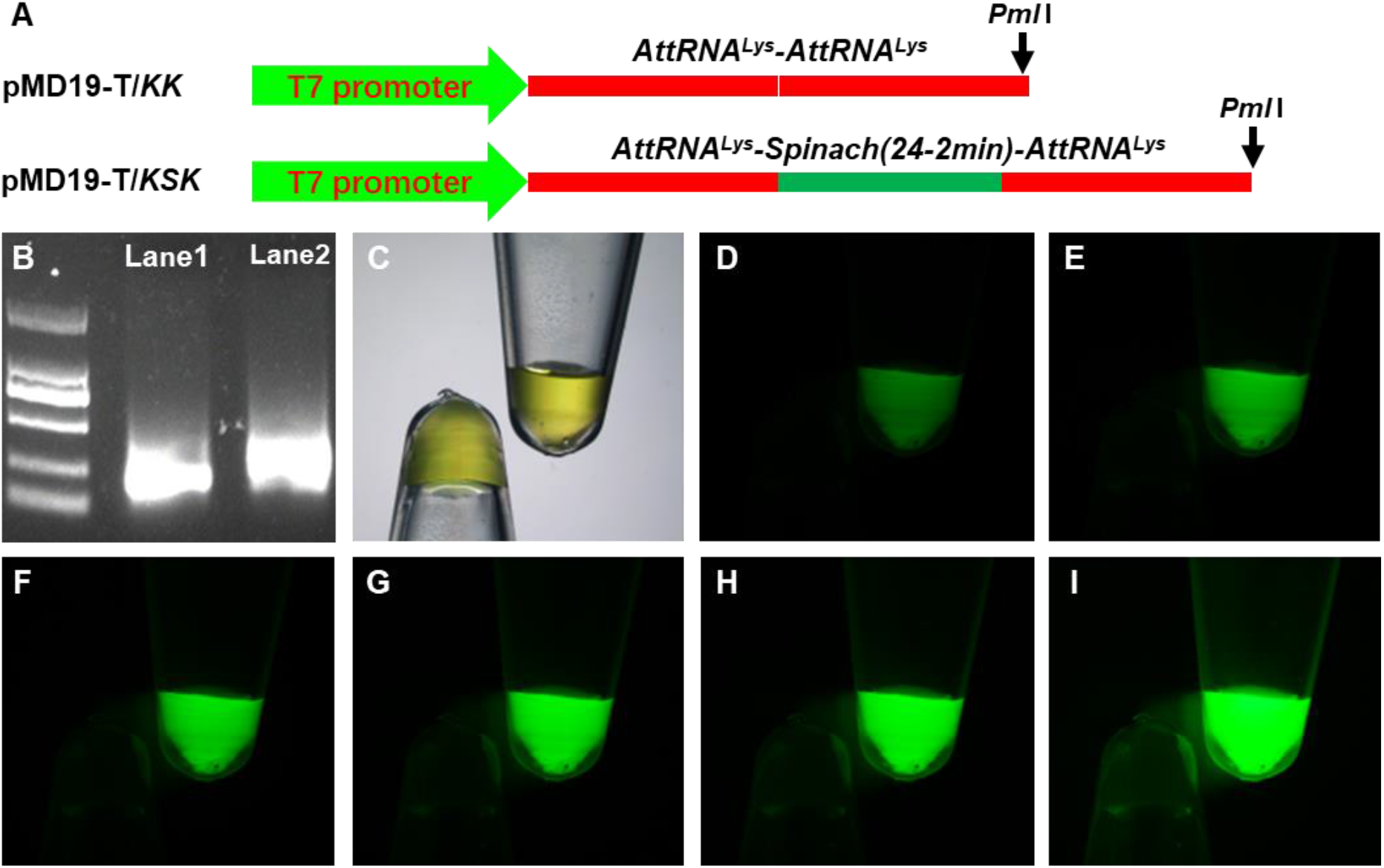
In vitro Spinach RNA fluorescence. A, Diagrammatic of KK and KSK expression cassettes in pMD19-T. KK (negative control) and KSK were transcribed from *Pml*I-linearized pMD19-T/KK or pMD19-T/KSK under the control of the T7 promoter. Detailed sequences of KK and KSK are included in Supplemental Data Set S1. B, 1.5% TAE-Agarose gel electrophoresis of KK and KSK RNA transcripts. KK and KSK RNA transcripts were loaded on Lane 1 and Lane 2, respectively. Marker: DM2000. C-I, KK and KSK RNA transcripts in 100 μM DFHBI solution. Photographs were taken under transmitted white light channel (C), or under FITC channel at the exposure time of 2 (D), 4 (E), 6 (F), 8 (G), 10 (H) and 20 (I) seconds using a Fluorescence Stereomicroscope. The concentration of KK and KSK RNA transcripts (C-I) was 2,915.70 and 2,969.55 ng/μl, respectively.

### Spinach-based RMG in onion epidermal cells and rice protoplasts

To transiently express Spinach in plant cells, we subcloned *KK* and *KSK* (Fig. 1A; Supplemental Data Set S1) into pEAQ-HT, a binary vector for efficient gene expression (Sainsbury and Lomonossoff, 2008; Sainsbury et al., 2009), and produced pEAQ-HT/KK and pEAQ-HT/KSK (Fig. 2A). Onion epidermal cells without chloroplasts were then bombarded with purified plasmid DNA of pEAQ-HT/KK or pEAQ-HT/KSK. We also bombarded onion tissues with pEAQ-HT/GFP (Sainsbury and Lomonossoff, 2008) to express GFP as positive control. GFP fluorescence was readily visible under the confocal microscope in onion epidermal cells 10-hr after bombardment (Fig. 2B and C). In striking contrast to KK control (Fig. 2D and E), strong green fluorescence was observed in onion epidermal cells that transiently expressed the KSK RNA in the presence of DFHBI (Fig. 2F and G; Supplemental Fig. S1). However, rice protoplasts (isolated from rice etiolated seedlings) expressing KSK or KK RNA under the control of the strong OsU6 promoter (Feng et al., 2013; Ma et al., 2015) appeared to have a similar level of green fluorescence in both cases (Supplemental Fig. S2A-D).

**Figure 2.**
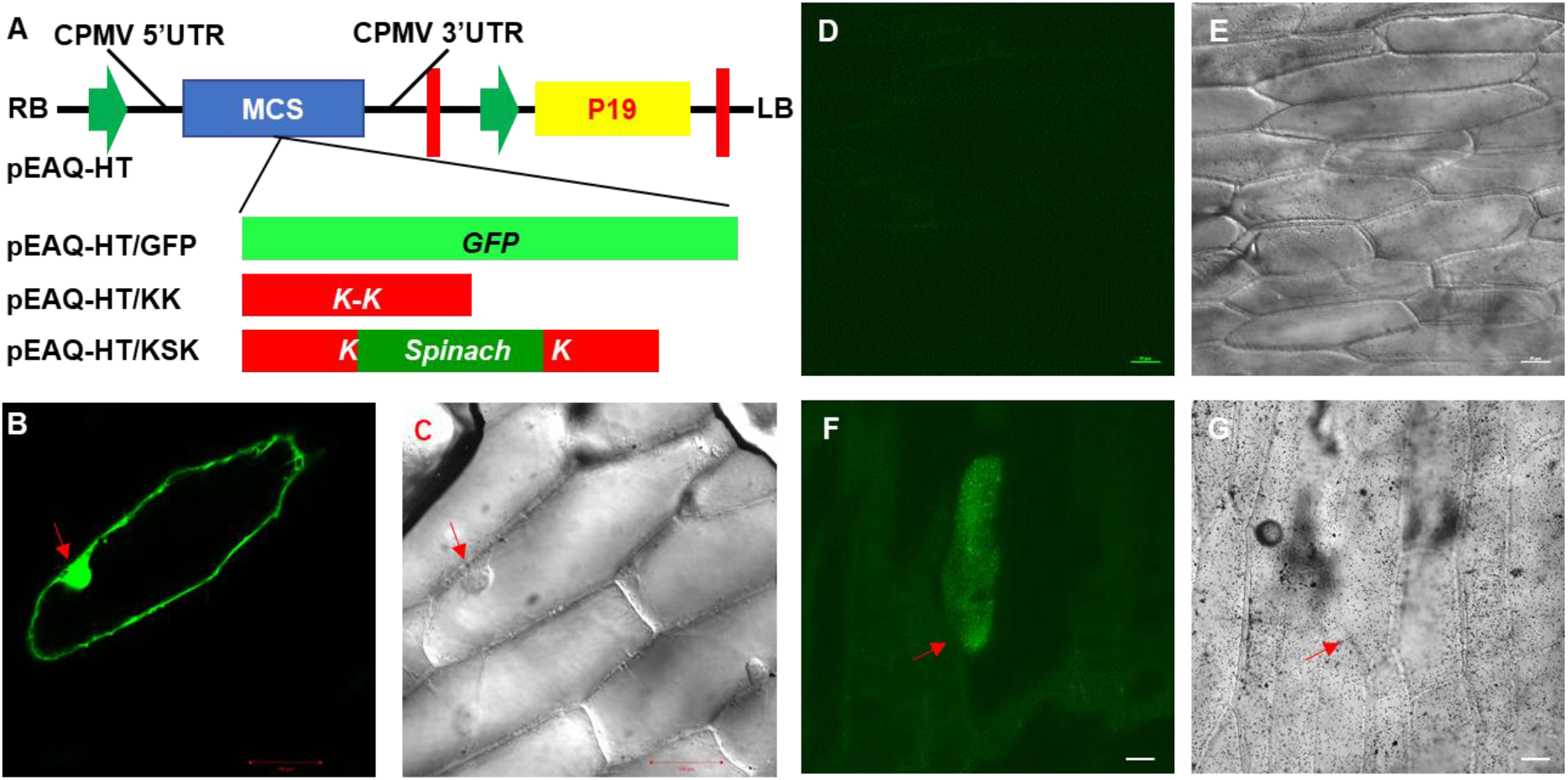
RMG in onion epidermal cells. A, Schematic of GFP, KK and KSK expression cassettes in pEAQ-HT. GFP, KK and KSK coding sequences were cloned into MCS of pEAQ-HT. Green arrows: 35S CaMV promoter sequences. Red thick lines: CaMV terminator sequences. MCS: multiple cloning site. CPMV 5’- and 3’-UTR: cowpea mosaic virus 5’ and 3’ untranslated regions which act as translational enhancer. P19: Tombusvirus silencing suppressor protein. B-G, RMG in onion epidermal cells. As a control, onion epidermal cells were bombarded with pEAQ-HT/GFP and showed GFP fluorescence at 12 hours after bombardment (HAB, B and C). Onion epidermis was bombarded with pEAQ-HT/KK (D and E) or pEAQ-HT/KSK (F and G). Green fluorescence was observed only in onion epidermal cells expressing KSK from with pEAQ-HT/KSK (F) at 12 HAB. Photographs were taken under FITC channel (B, D and F) or through transmitted light (C, E and G). Bar = 100 μm in B and C; bar = 50 μm in D-G; Red arrows indicate cells showing green fluorescence.

### Transgenic expression of Spinach in *Nicotiana benthamiana*

To obtain stable Spinach expression, we transformed *N. benthamiana* with pEAQ-HT/KK and pEAQ-HT/KSK and generated two independent transgenic lines with a single copy of a transgene for constitutive expression of KK or KSK RNA (Fig. 3). The insertion of the *KK* and *KSK* transgenes in the *N. benthamiana* genome was readily detected by genomic PCR (Fig. 3A). Moreover, RT-PCR assays revealed that KK and KSK RNAs were being expressed in *KK* and *KSK* transgenic plants, respectively (Fig. 3B). However, the level of green fluorescence seen in total RNAs extracted from KK and KSK RNA-expressing transgenic plant leaf tissues in the presence of the DFHBI fluorophore were of similar intensity (Fig. 3C and D; Supplemental Fig. S3; S4 and S5). Direct visualization of the cryo-sectioned leaf tissues or protoplasts derived from the *KK* and *KSK* transgenic plants also showed no noticeable difference in green fluorescence with or without DFHBI (Fig. 3E-H). These results indicate that stable expression of *KSK* RNA from a single locus in transgenic plants was insufficient for *in vivo* monitoring of cellular spinach RNAs in plants.

**Figure 3.**
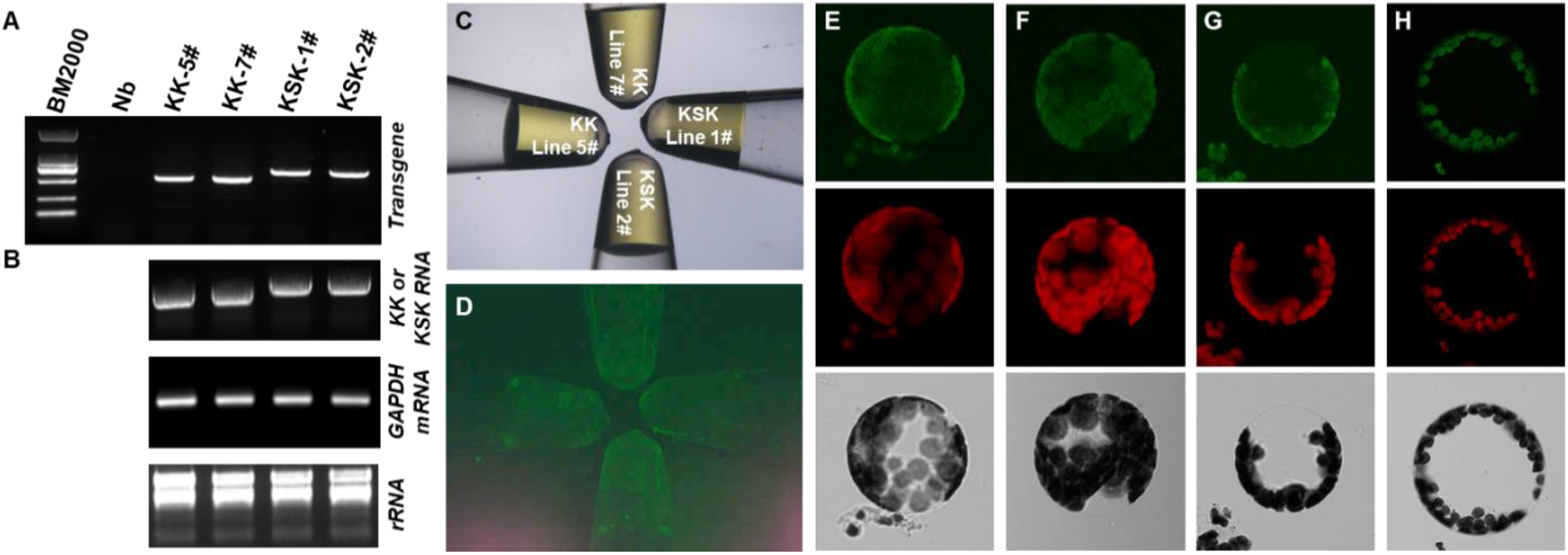
RMG in transgenic plants. A, Genomic PCR confirmation of *KK* and *KSK* transgenes in independent *Nicotiana benthamiana* transformants. B, Transgenic expression of *KK* and *KSK*. RT-PCR detection of KK and KSK RNA transcripts (Top panel), and *GAPDH* mRNA (Middle panel). PCR consisted of 25 cycles for GAPDH or 35 cycles for *KK* and *KSK*. Total RNA was also analyzed by 1% agarose-TAE gel electrophoresis (Bottom panel). C and D, Ex vivo fluorescence of plant RNAs. Twenty microliter solution with 100-μM DFHBI and 750-μg/μl total RNA extracted from young leaf tissues of the *KK* and *KSK* transgenic plants was photographed under transmitted white light (C) or FITC filter (D). E – H, RMG in protoplasts. Protoplasts were isolated from the *KK* (E and F) or *KSK* (G and H) transgenic plant leaves with (F and H) or without (E and G) 400-μM DFHBI. Photographs were taken under FITC (Top panel), red (Middle panel) filter or transmitted light (Bottom panel).

### Delivery of Spinach to plants via virus expression systems

The failure of stable expression of KSK RNA to distinguish fluorescent cells and tissues may be due to the low level of the KSK RNA produced from a single copy of the transgene in the transgenic plants. To overcome this potential issue, we opted to try plant virus-based expression systems as these often produce high amount of RNAs (and proteins) from recombinant viral RNA or DNA genomes during virus infection of plants (Qin et al., 2015; Qin et al., 2017; Lai et al., 2020). We cloned *KK* and *KSK* into a Potato virus X (PVX)-based vector (van Wezel et al., 2001) and produced PVX/KK and PVX/KSK (Fig. 4A; Supplemental Data Set S1). Three additional constructs PVX/AtFT:KSK, PVX/mAtFT:KSK and PVX/AtTFL1:KSK were also included in this study (Fig. 4A). These constructs were expected to express wild-type and mutant Arabidopsis *Flowering Locus T(AtFT* and *mAtFT*; Li et al., 2009; Li et al., 2011; Lu et al., 2012; Ellison et al., 2020) mobile RNA or the Arabidopsis *Terminal Flowering* (*AtTFL1*; Lu et al., 2012) immobile RNA, all tagged with the KSK aptamer.

**Figure 4.**
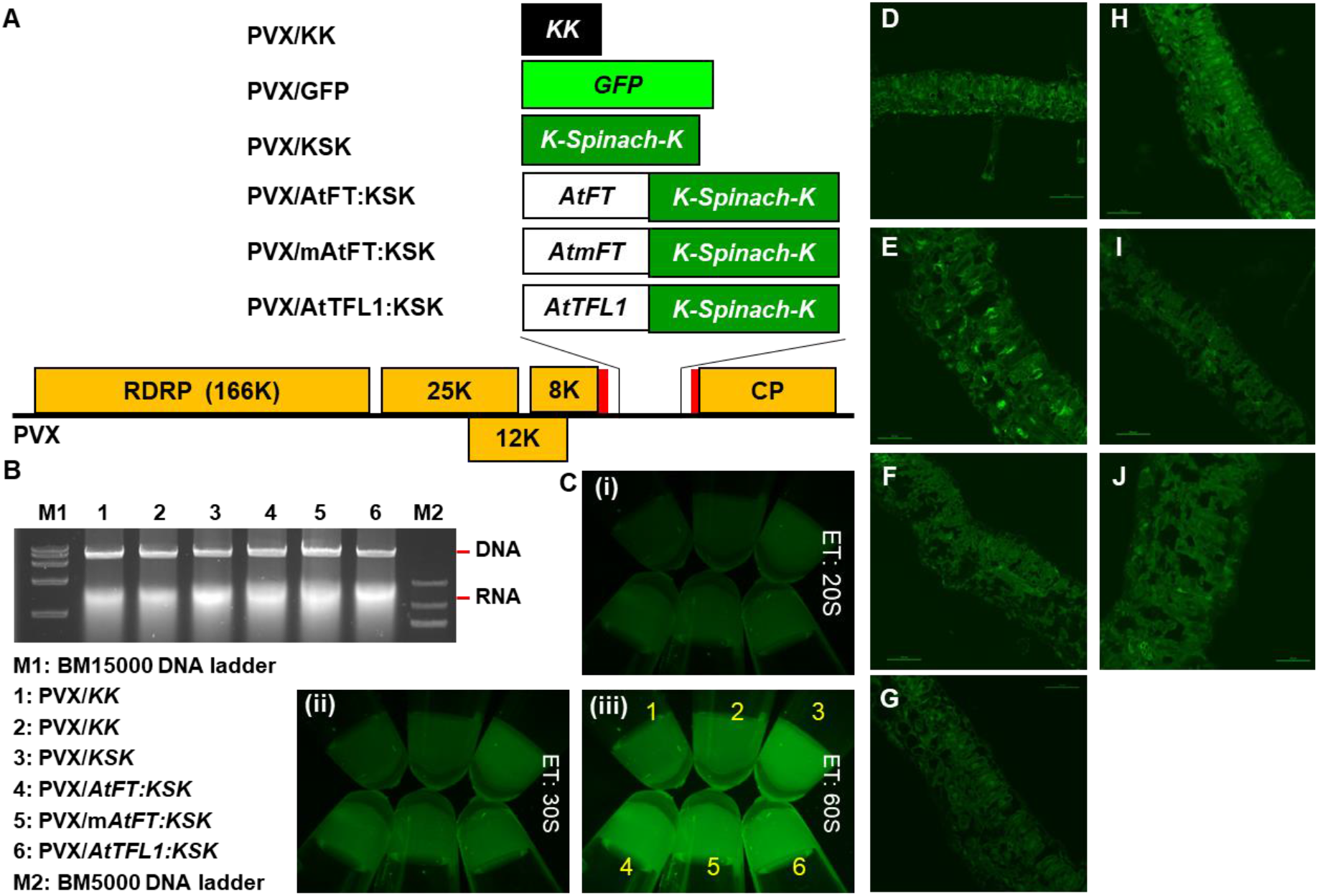
Virus-based RMG. A, Potato virus X (PVX)-based RMG constructs. The RNA genome of PVX encodes an RNA-dependent RNA polymerase (RDRP), a triple-gene block for three movement proteins of 25K, 12K and 8K, as well as a coat protein (CP). GFP, KK, KSK and the wild-type and mutant Arabidopsis *FT* and *TFL1* genes were cloned into the PVX-based vector as outlined. The two thin red boxes indicate the duplicated CP subgenomic RNA promoter. B, Recombinant PVX RNA transcripts. Transcripts were generated by in vitro transcription and analyzed through a 1% Agarose-TAE gel. C, Fluorescence of recombinant PVX RNAs generated by in vitro transcription. The number corresponding to in vitro RNA samples (10 μg) are same as indicated in B. Exposure time (ET) for photographing was 20 (i), 30 (ii), and 60 (iii) seconds (S). D-J, Fluorescent images of cryo-sections of systematic leaves. *N. benthamiana* plants were mock-inoculated (D), or infected with PVX/GFP (E); PVX/KK (F); G. PVX/KSK (G); H. PVX/AtFT:KSK (H); I. PVX/AtmFT:KSK (I); or PVX/AtTFL1:KSK (J). Sections were immersed in 200 μM DFHBI. D-J. Bar = 50 μM (D-J).

Recombinant PVX RNAs were sufficiently generated by *in vitro* transcription (Fig. 4B). Indeed, these viral recombinant RNAs with the KSK RNA tag (PVX-KSK, PVX-(m)AtFT-KSK or PVX-AtTFL1-KSK RNA) emitted stronger green fluorescence than the PVX-KK RNA (Fig. 4C). However, the fluorescent intensity was much weaker than free KSK RNA (Fig. 1). These results suggest that embedding *KSK* in the PVX genome (of approximately 6.5 kilobases in size) may attenuate the capacity of KSK RNA binding to DFHBI for emitting green fluorescence. Nevertheless, we infected wild-type *N. benthamiana* plants with each of the recombinant PVXs or PVX/GFP (Fig. 4A). Systemic young leaf tissues with chlorosis systems, characteristic of PVX infection, were cryo-sectioned and immersed with a DFHBI solution. We observed green fluorescence in tissues infected with PVX/KSK, PVX/AtFT:KSK, PVX/mAtFT:KSK, PVX/AtTFL1:KSK; however, the fluorescence was much weaker than that observed in tissues infected with PVX/GFP, and often indistinguishable from PVX/KK infected samples (Fig. 4D-J). A similar result was also obtained using a geminivirus (DNA virus)-based expression vector (Supplemental Fig. S6; Tang et al., 2010). Taken together, our results indicate that, as with stable transgenic expression, a monomer of KSK RNA delivered from two highly efficient virus expression vectors was inadequate to visualize RNAs in plant cells and tissues.

### Direct and indirect assays of RNAs tagged with tandem repeats of Spinach

The fluorescent signal emitted from a monomer of Spinach from both transgenic and viral transient expression systems was too weak to be differentiated from background noise fluorescence in plants. To enhance the “signal vs noise” ratio, we cloned a tandem repeat of 2 to 5 spinach aptamers into the PVX vector, as described in a recent report (Zhang et al., 2015), and generated PVX/KSK*2, PVX/KSK*3, PVX/KSK*4 and PVX/KSK*5 (Fig. 5A). We also renamed PVX/KSK (Fig. 4A) as PVX/KSK*1 hereafter. Indeed, recombinant viral RNA PVX-KSK*1, PVX-KSK*2, PVX-KSK*3, PVX-KSK*4 and PVX-KSK*5 showed an increasing capacity to emit green fluorescence in the presence of DFHBI (Fig. 5B-F). We also noticed that PVX-KK RNA-DFHBI mixed solution possessed some background fluorescence (Fig. 5F). Using the equation drawn from the FITC vs fluorescent intensity standard curve (Supplemental Fig. S4A-C and Supplemental Fig. S5), the concentration equivalent to FITC was calculated to be 0.0474 μM for in vitro KSK*5 RNA transcripts after deducting the background of the KK DFHBI fluorescence (Fig. 5F).

**Figure 5.**
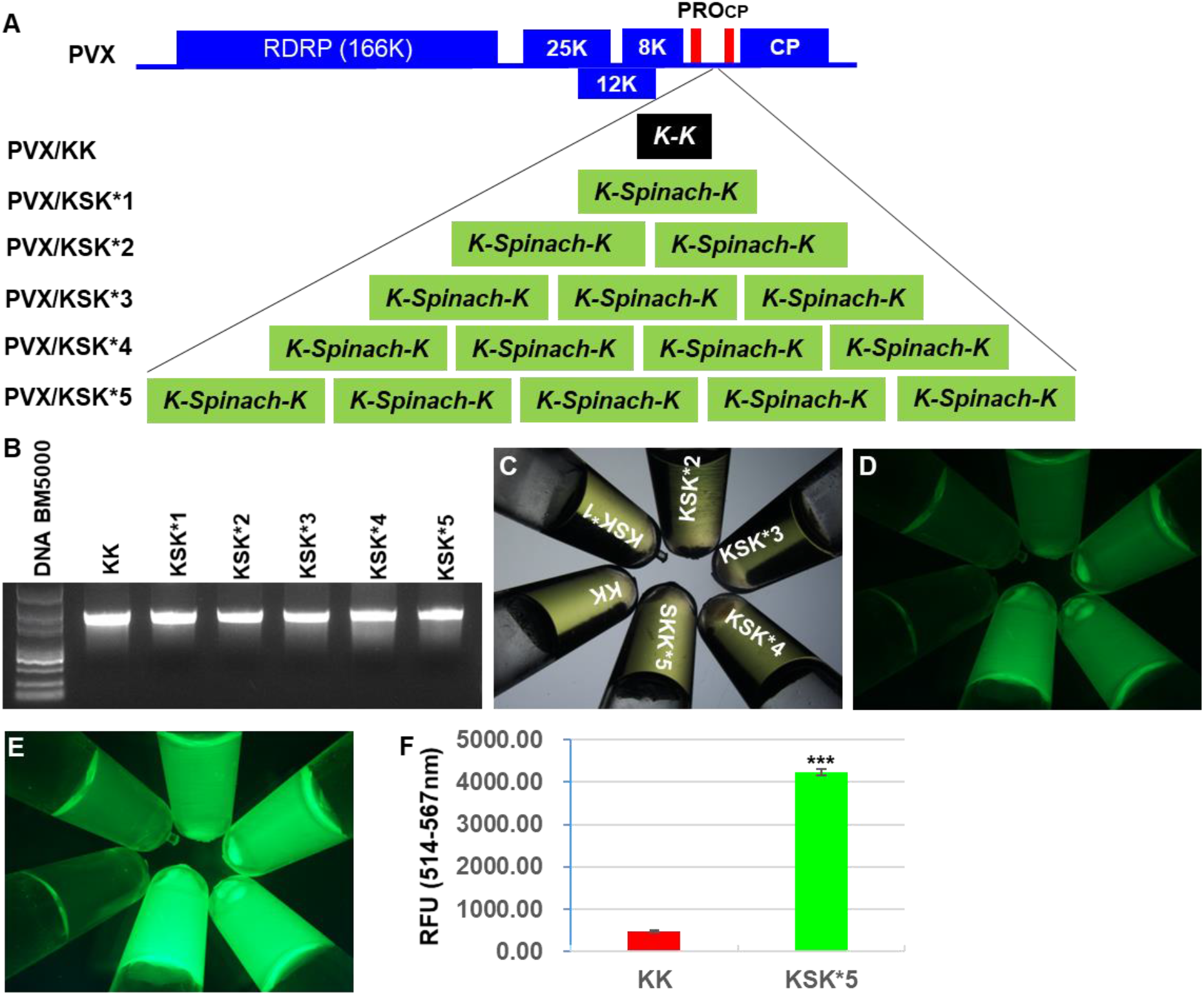
Tandem repeats of Spinach enhance fluorescence in vitro. A, Schematic diagrammatic of the tandem repeats of spinach in PVX-based vector. The RNA genome of PVX encodes an RNA-dependent RNA polymerase (RDRP), a triple-gene block for three movement proteins of 25K, 12K and 8K, as well as a coat protein (CP). Expression of KK, KSK or repeated KSK RNA was under the control of the duplicated CP subgenomic RNA promoter (PRO_CP_, thin red box). B, Production of KK and KSK RNA transcripts. 1 μg in vitro recombinant PVX RNA was analyzed by 1% Agarose-TAE gel electrophoresis. C-E, In vitro fluorescence of PVX recombinant RNA. PVX recombinant RNA samples (10 μg) containing 200uM DFHBI were observed through transmission detector (C) and FITC filter (D and E). Exposure time was 40 (D) and 150 (E) seconds, respectively. The order of samples is the same as indicated in C. F, Quantitative fluorescence. RNA fluorescence was measured as relative fluorescence units (RFU) for PVX/KK and PVX/KSK*5 transcripts generated by in vitro transcription. The RFU value is shown as Mean ± SD (n = 3). Student’s ***t***-test showed a significant difference between PVX/KK and PVX/KSK*5 (P ≤ 3.05 x 10^−7^, indicated by 3 asterisks). Using the equation drawn from the FITC vs fluorescence standard curve (Supplemental Fig. S3A-C and Supplemental Fig. S4), we calculated the fluorescence emitted from the ‘in vitro RNA transcripts’ for PVX/KK and PVX/KSK*5 was equivalent to that emitted by 0.1044 μM and 0.1518 μM FITC respectively.

We then inoculated young leaves of wild-type *N. benthamiana* with recombinant viral RNA PVX-KK, PVX-KSK*1, PVX-KSK*2, PVX-KSK*3, PVX-KSK*4 or PVX-KSK*5. These plants became systemically infected and developed clear chlorosis on newly growing young leaves at 7 days post-inoculation (DPI) and onwards (Supplemental Fig. S7A-F). At 17 DPI, total RNA was extracted from young leaf tissues and mixed with DFHBI. We observed stronger ex vivo fluorescence from RNA extracted from tissues infected with PVX/KSK*5 than PVX/KK (Fig. 6A-F). The relative increase in fluorescent intensity of total RNA isolated from plants infected with PVX/KSK*1, PVX/KSK*2, PVX/KSK*3, PVX/KSK*4 and PVX/KSK*5 was very similar to that observed for *in vitro* produced RNA transcripts from these constructs (Fig. 5C; Fig. 6A-F). Similarly, we also detected some background ex vivo fluorescence from total RNA extracted from PVX/KK-infected leaf tissues (Fig. 6F). Upon calculation using the equation based on the FITC vs fluorescent intensity standard curve (Supplemental Fig. S4A-C and Supplemental Fig. S5), the concentration equivalent to FITC was 0.0221 μM for total RNAs extracted from PVX/KSK*5-infected leaf tissues after deducting the background KK DFHBI fluorescence (Fig. 6F). Despite the positive observations of ex vivo RMG, RNA fluorescence in cryo-sections of young leaves infected with PVX/KK (Fig. 6G) or PVX/KSK*5 (Fig. 6H) was not sufficiently different.

**Figure 6.**
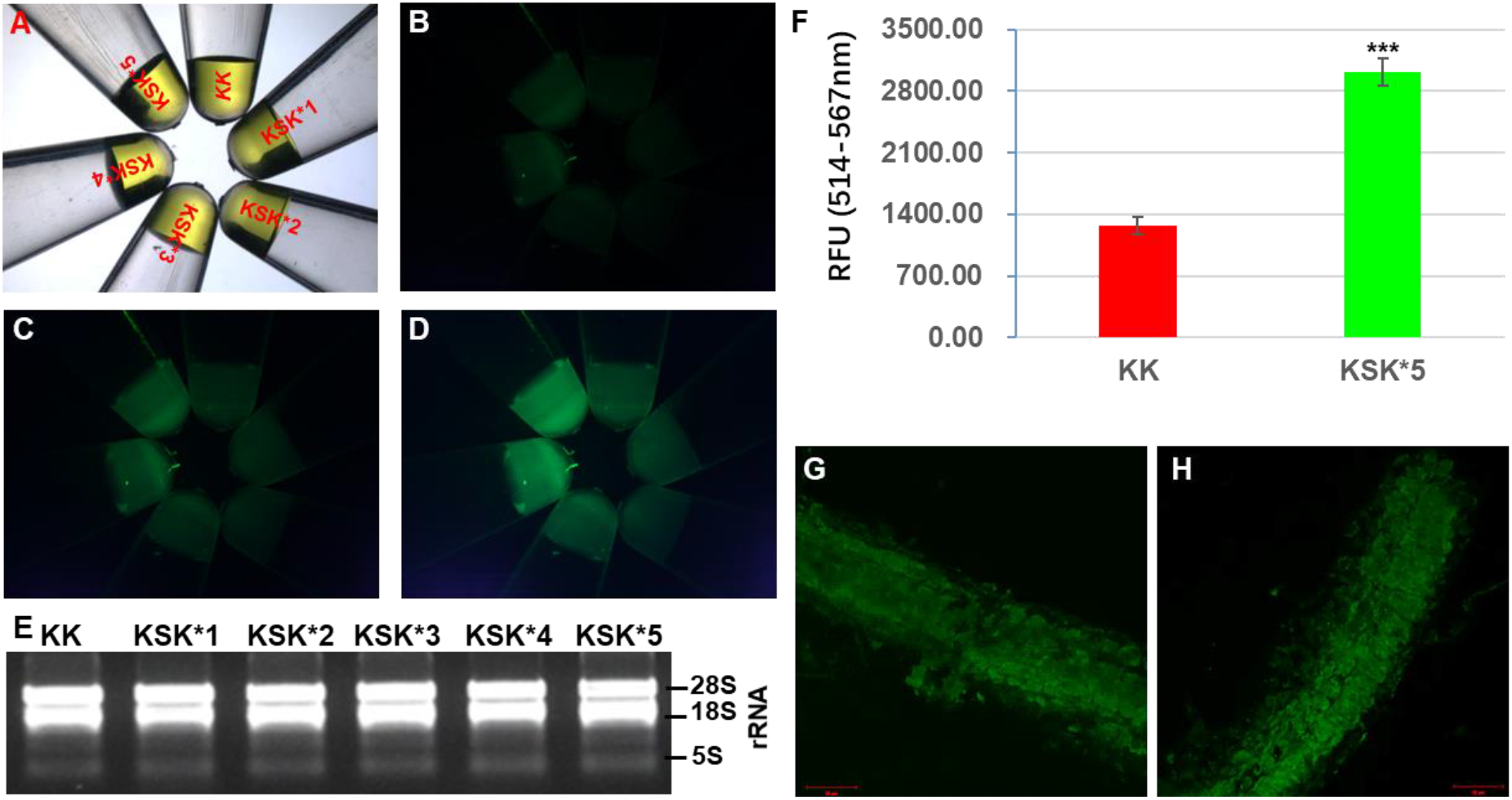
PVX/KSK*5-mediated RMG. A-D, Ex vivo analysis of plant RNAs. 10 mg of total RNA extracted from *N. benthamiana* young leaf tissues infected with PVX/KK (KK), PVX/KSK*1 (KSK*1), PVX/KSK*2 (KSK*2), PVX/KSK*3 (KSK*3), PVX/KSK*4 (KSK*4) or PVX/KSK*5 (KSK*5) were mixed with 1-mM DFHBI and photographed under transmitted light (A) or FITC channel (B-D). Exposure time was 150 (B), 300 (C) and 600 (D) seconds, respectively. E, Gel analysis of total plant RNA. 5S, 18S and 28S rRNA are indicated. F, Quantitative fluorescence. RNA fluorescence was measured as relative fluorescence units (RFU) for total RNA extracted from plant leaves infected with PVX/KK and PVX/KSK*5. The RFU value is shown as Mean ± SD (n = 3). Student’s ***t***-test showed a significant difference between PVX/KK and PVX/KSK*5 (P ≤ 0.00035, indicated by 3 asterisks). G and H, Fluorescent images of cryo-sections of young systematic leaves of N. benthamiana plants infected with PVX/KK (G) or PVX/KSK*5 (H). Sections were treated with 200-μM DFHBI. Using the equation drawn from the FITC vs fluorescence standard curve (Supplemental Fig. S3A-C and Supplemental Fig. S4), we calculated the fluorescence emitted from total RNA extracted from PVX/KK and PVX/KSK*5-infected leaf tissues was equivalent to that emitted by 0.1143 μM and 0.1364 μM FITC respectively.

## DISCUSSION

Direct visualization of RNA has attracted a great deal of interest in plant science, particularly in RNA metabolism and mobile RNA signaling. Unfortunately, the successful establishment of Spinach RNA-mimicking GFP in prokaryotic and eukaryotic cells some ten years ago (Paige et al., 2011) has not led to establish a similar technology in plants. The only attempt to use the Spinach aptamer to monitor plant cellular RNAs was unsuccessful (Huang et al., 2017). In this study, we re-evaluated the use of Spinach-based RNA visualization in plants via a range of different expression strategies. Our pump-priming work showed this technology could be used for ex vivo and in vivo monitoring of RNAs in onion and *N. benthamiana*.

We showed that, similar to bacterial, yeast and human cellular tRNAs, plant tRNAs such as tRNA^Lys^ can protect and/or stabilize spinach RNA aptamer enabling interaction with the DFHBI fluorophore and emission of sufficient green fluorescence for in vitro detection (Fig. 1). We also observed that tagging spinach RNA to longer RNAs such as the PVX genomic RNA could lead to attenuation of the spinach fluorescence signal (Fig. 3)

We also showed that, tRNA^Lys^-spinach-tRNA^Lys^ (KSK), once delivered into “chloroplast-free” onion epidermal cells via bombardment, can emit detectable green fluorescence in the presence of DFHBI (Fig. 2; Supplemental Fig. S1). However, in rice protoplasts (single cells) isolated from green-yellowing stem/leaf tissues, KSK-emitted fluorescence became less distinguishable from the KK control (Supplemental Fig. S2). This may be due to the background noise green fluorescence resulted from chromoplasts in rice protoplasts which were isolated from the etiolated stem tissues. Nevertheless, contrary to the previous report (Huang et al., 2017), our work clearly demonstrates that the spinach based RMG can work in plants, at least in onion epidermal cells.

Transgenic expression in stably transformed *N. benthamiana* or virus-based transient expression of monomer KSK in wild-type *N. benthamiana* plants, failed to generate clear evidence that KSK-DFHBI could emit distinguishable green fluorescence when compared to negative KK controls (Fig. 3; Fig. 4; Supplemental Fig. S3 and Supplemental Fig. S6). This was likely due to a high background fluorescence masking the genuine fluorescent signal emitted from KSK-DFHBI. The background fluorescence might come from either auto-fluorescence of cellular components such as chloroplasts and chromoplasts (as seen with the rice protoplasts (Supplemental Fig. S2)), or from DFHBI non-specific binding to cellular RNAs. The latter was indeed evidenced by our serendipitous finding that DFHBI could directly and specifically stain RNA, but not DNA in agarose gels; showing strong fluorescence under long-wavelength UV light (Supplemental Fig. S8). This result is also consistent with the original discovery in which the Spinach-specific fluorescence was often observed after the background noise fluorescence of negative controls was subtracted (Paige et al., 2011).

We developed the system further by incorporating tandem repeats of spinach RNA into recombinant PVX RNAs, and by doing this we were able to detect in vitro and ex vivo (ex planta) KSK-specific green fluorescence although in vivo (in planta) RMG was less apparent (Fig. 5 and Fig. 6).

In summary, we report that the Spinach RNA aptamer can mimic GFP in plant cells. However, this technology requires further improvement to realize its full potential in plant RNA visualization. The main challenge is to reduce the background fluorescence whilst enhancing KSK-specific green fluorescence. This could be achieved via expressing more tandem repeats of spinach, as demonstrated in this work, or its derivatives in plants. Indeed, since the development of Spinach (Paige et al., 2011), several new aptamers such as Spinach2 (Strack et al., 2013), Baby Spinach (Huang et al., 2014), iSpinach (Autour et al., 2016), Pandan (Aw et al., 2016), Broccoli (Filonov et al., 2014), RNA-Mango (Dolgosheina et al., 2014), Corn-DFHO (Warner et al., 2017) and Pepper (Chen et al., 2019) have been uncovered and used for RNA visualization. More recently, a series of fluorescent aptamers based on the modified three-way junction scaffold and the optimized Broccoli has been reported to be able to visualize RNA in plants (Bai et al., 2020). Moreover, Spinach2, Baby Spinach and iSpinach also use DFHBI as fluorophore, and these derivatives are superior to Spinach in terms of luminescence intensity and/or the light quenching property (Filonov et al., 2014; Warner et al., 2017; Chen et al., 2019). These newly developed aptamers do not require any tRNA scaffolds for protection and they can still stably bind to RNA and stimulate fluorescence (Filonov et al., 2014; Chen et al., 2019). This can be beneficial for the virus-based delivery system because recombinant plant viruses tend to lose larger insert from their genomes while infecting plants (Qin et al., 2015). Thus, these newer RNA aptamers offer more options for RNA fluorescence *in planta*. An alternative strategy to reduce fluorescent noise is to screen novel fluorophore(s) that may be more specific than DFHBI in their binding to Spinach (Supplemental Fig. S8). Such a fluorophore(s), if identified, will certainly be useful to enhance the effectiveness of Spinach-based RMG in plants.

## MATERIALS AND METHODS

### Plant Materials and Growth

*Nicotiana benthamiana* plants were grown in insect-free growth room at 25°C during day and 18°C at night under a 50% humidity environment with a 16-hr day/8-hr night periodic cycle.

### Construction of Vectors

Original sequences including (i) 73-nucleotides (nt) *AttRNA^Lys^* (K), (ii) 80-nt *Spinach* (S), (iii) 152-nt *AttRNA^Lys^-AttRNA^Lys^* (KK), (iv) 250-nt *AttRNA^Lys^-Spinach-AttRNA^Lys^* (KSK), (v) 227-nt T7 promoter-KK, (vi) 374-nt T7 promoter-K-Spinach-K (KSK), (vii) 205-nt pCVA-KK, and (viii) 352-nt pCVA-KSK are listed in Supplemental Data Set S1. To obtain double-stranded (ds) KK DNA fragment, a pair of oligonucleotides P001 and P002 (Supplemental Table S1) were annealed to form a dsDNA molecule. Then a second pair of oligonucleotides P003 and P004 (Supplemental Table S1) were also annealed together. The two dsDNA fragments were cloned into the *Mlu*I/*BspE*I sites of the *Potato virus X*(PVX) based vector (van Wezel et al., 2001) to generate PVX/KK. An *Eag*I site was introduced between the two Ks (Fig. 4A). The KK fragment was then amplified from PVX/KK using different sets of primers (Supplemental Table S1) and subcloned into pMD19-T (TAKARA), the *Bbs*I site of pBluescript SK+/OsU6 (Feng et al. 2013), the *Nru*I/*Xho*I sites of pEAQ-HT (Sainsbury et al., 2009) or the *Xba*I/*Kpn*I sites of pCVA (Chen et al., 2015) to produce pMD19-T/KK (Fig. 1A), the pBluescript SK+/KK (Supplemental Fig. S2A), pEAQ-HT/KK (Fig. 2A) and pCVA/KK (Supplemental Fig. S6A), respectively. The T7 promoter sequence and a unique *Pml*I site were introduced to the 5’- or 3’-end of KK in pMD19-T/KK, respectively (Fig. 1A).

We cloned the KSK dsDNA fragment which was commercially produced by Invitrogen into the *Age*I/*Sma*I sites of pEAQ-HT and generated pEAQ-HT/KSK (Fig. 2A). The KSK fragment was then amplified from pEAQ-HT/KSK using different sets of primers (Supplemental Table S1) and subcloned into pMD19-T, the *Bbs*I site of pBluescript SK+/OsU6, the *Mlu*I/*Eag*I or *Eag*I/*Sal*I sites of PVX, or the *Xba*I/*Kpn*I sites of pCVA to produce pMD19-T/KSK (Fig. 1A), the pBluescript SK+/KSK (Supplemental Fig. S2A), PVX/KSK(1) and PVX/KSK(2) (Fig. 4A; Fig. 5A), and pCVA/KSK (Supplemental Fig. S6A), respectively. For pMD19-T/KSK, T7 promoter sequence was incorporated at the 5’-end of KSK while a *Pml*I site was introduced at the 3’ end of KSK (Fig. 1A).

Construction of PVX/AtFT:KSK, PVX/mAtFT:KSK and PVX/AtTFL1:KSK were all based on PVX/KSK(2) in which KSK was cloned into the *Eag*I/*Sal*I sites (see above). Wild-type and non-sense mutant Arabidopsis *FT* genes *AtFT* and *mAtFT* were amplified (Li et al., 2009) and cloned into the *Cla*I/*Eag*I sites of PVX/KSK(2) to generate PVX/AtFT:KSK and PVX/mAtFT:KSK (Fig. 4A). Similarly, the AtTFL1 gene was amplified (Li et al., 2011) and cloned into *Cla*I/*Mlu*I sites of PVX/KSK(2) to generate PVX/AtTFL1:KSK (Fig. 4A).

Tandem repeats of KSK were constructed in PVX/KSK(1) in which KSK was cloned into the *Mlu*I/*Eag*I sites of the PVX vector (see above). Insertion of an extra KSK into the *Eag*I/*BspE*I sites of PVX/KSK(1) (aka PVX/KSK*1) produced PVX/KSK*2, then a KSK into the *EcoRV*/*Sal*I sites of PVX/KSK*2 brought about PVX/KSK*3. To construct the PVX/KSK*4, a KSK was cloned into the *BspEI/EcoRV* sites of PVX/KSK*3; and PVX/KSK*5 was constructed by inserting a KSK into the *Cla*I/*Mlu*I sites of PVX/KSK*4 (Fig. 5A).

The integrity of the sequence insertions in all constructs was confirmed by Sanger sequencing.

### Preparation of DFHBI Solution

Fluorophore DFHBI (3,5-difluoro-4-hydroxybenzylidene imidazolinone) was bought from Lucerna™ company (http://www.lucernatechnologies.com/fluorophores-c17/). DFHBI was dissolved in DMSO to prepare a 40 mM stock solution. It was then diluted with 100 mM HEPES buffer (pH 7.5) to produce a 2 mM DFHBI/5% DMSO working solution (Paige et al., 2011). In this work, the final concentration of DFHBI used to trigger spinach fluorescence was 100-200 μM for spinach-tagged RNA transcripts generated by in vitro transcription, or 1-2 mM for total RNAs extracted from plant tissues and for in planta RNA mimicking GFP (RMG) assay.

### Particle Bombardment and Confocal Microscopy

Plasmid DNA of pEAQ-HT/KK and pEAQ-HT/KSK was prepared from *Escherichia coli* 2 T1^R^ cells (Thermo Fisher Scientific) using QIAprep Spin Miniprep Kit, and their concentration was adjusted to 1μg/μl. Gold particles were coated with DNA and onion epidermal cells were particle-bombarded as described (Wydro et al., 2006; Ding et al., 2009). Briefly, 1.5 mg of gold microcarriers (1 μm in diameter) were washed with 70% ethanol once and then 100% ethanol twice. After a quick spin, the clean gold microcarriers were collected, air-dried and resuspended in 50μl 50% glycerol. Then 10μg plasmid DNA, 50μl 2.5M CaCl_2_, 20μl 0.1M spermidine and 250μl 70% ethanol were mixed sequentially and progressively. After a vigorous vortex for 2-3 seconds, followed by a quick spin, the DNA-coated gold microcarriers were collected, air-dried and resuspended in 30μl 100% ethanol. 10 μl DNA-coated gold microcarriers were dropped onto a microparticles carrier disk (Macrocarriers #1652335, Bio-Rad) and bombardment was carried out using a PDS-1000/He Biolistic Particle Delivery System (Bio-Rad). After 12 hours culture in a hypertonic medium (0.8 % Phytagel half-strength Murashige and Skoog (MS) basal medium, 0.256 M (46.67 g/L) sorbitol and 0.256 M (46.67 g/L) mannitol), onion epidermis was immersed into 100 μM DFHBI (3,5-difluoro-4-hydroxybenzylidene imidazolinone) for 30min, examined and photographed using a Zeiss LSM 710 confocal laser scanning microscope (Ding et al., 2009; Paige et al., 2011).

### Plant Transformation

*N. benthamiana* was transformed by a leaf disc procedure with *Agrobacterium tumefaciens* GV3101 harboring pEAQ-HT/KK or pEAQ-HT/KSK (Hong et al., 1996; Sainsbury et al., 2009). Primary transformants were selected for resistance to 50 μg/ml kanamycin and further verified by genomic PCR. Following self-pollination, segregation of T1 and T2 progenies was tested for sensitivity to 50 μg/ml kanamycin. Two homozygous lines with a single of copy of the transgene (as indicated by a 3:1 segregation ratio) for KK (Line 5 and Line 7) or KSK (Line 1 and Line 2) were selected for RMG analysis.

### Agroinfiltration Assay

Agroinfiltration assay was performed in young leaves of *N. benthamiana* as previously described (Bruce et al., 2011; Yu et al., 2018). Briefly, 10 ml of A. *tumefaciens* GV3101 carrying pCVA/KK, pCVA/KSK or pCVB, a *Cabbage leaf-curl geminivirus* (CLCV)-based vector (Tang et al., 2010; Chen et al., 2015) were cultured to 0.6-0.8 OD_600_. Agrobacterium was then collected by centrifugation at 4,000 rpm for 10 min, and re-suspended to make the final density of OD_600_=2.0 in sterilized water. Agrobacterium with pCVA/KK or pCVA/KSK was mixed with an equal volume of agrobacterium with pCVB and infiltrated to young leaves of *N. benthamiana* plants at the six-leaf-stage through a needleless syringe. At 3 day-post-agroinfiltration, leaf tissues were cryo-sectioned, stained with 200μM DFHBI, examined and photographed using a Zeiss LSM 710 confocal laser scanning microscope. Five milligrams of total RNAs extracted from agroinfiltrated tissues in 1 mM DFHBI solution were examined and photographed using a Nikon fluorescent stereomicroscope (SMZ1500) equipped with a DN100 Digital Internet Camera (Nikon) through a white-light or FITC (fluorescein isothiocyanate) filter. Exposure time was up to 600 seconds dependent on the FITC fluorescent intensity of samples.

### Cryo-Sectioning

Plant tissue fixation, embedding and cryo-sectioning were carried out as described (Dong et al., 2003). Briefly, young leaves from *KK* or *KSK* transgenic plants or virus-infected *N. benthamiana* leaves were cut into 3 – 5-mm wide strips. Those small leaf tissues were embedded in the Tissue-Tek (OCT Compound Torrance) and sectioned in a cryostat at −25°C (Bright Instruments OTS). Ten to fifty-micrometer sections were mounted in 50% glycerol onto RNase-free slides (Thermo Fisher Scientific). After adding 20μl 200μM DFHB, sections were covered by RNase free cover glasses (Thermo Fisher Scientific), examined and photographed using a Zeiss LSM 710 confocal laser scanning microscope.

### Protoplast Preparation

Rice (*Oryza sativa* L. cv. Nipponbare) grains were sterilized and sowed on 8% agar rice medium (Yoshida et al., 1976) and cultured in dark at 28 °C for 10-12 days. Stems of etiolated rice seedlings were cut into 0.5-mm pieces by a sharp blade. The small stem slices were then immersed in 0.6 M mannitol for 10 minutes (Chen et al., 2006). On the other hand, young leaves of homozygous *KK* or *KSK* transgenic *N. benthamiana* lines were collected from one-month old plantlets grown in half-strength MS agar medium, and cut into 3 by 6-mm wide-length strips by a sharp blade. The small rice stem pieces and the *KK or KSK* transgenic leaf strips were digested in W5 solution (2 mM MES, 154 mM NaCl, 125 mM CaCl_2_ and 5 mM KCl, pH=5.7) with 1.5% cellulase R10 and 0.4% macerozyme R10 for 2-3 hours at room temperature in dark to release protoplasts. Protoplasts were collected by centrifugation at 100 g after filtrating out undigested stem or leaf tissues through 75 μm nylon mesh, and re-suspended in MMG solution (4 mM MES, 0.4 M mannitol and 15 mM MgCl2, pH=5.7) with 1 mM DFHBI (Yoo et al., 2007). *KK* or *KSK* transgenic *N. benthamiana* protoplasts were examined and photographed using a Zeiss LSM 710 confocal laser scanning microscope.

### PEG-mediated Transfection

Ten micrograms of pBluescriptSK+/KK or pBluescriptSK+/KSK plasmid DNA were mixed gently with 100 μl rice protoplast resuspension (2×10^4^). To the protoplast and DNA mixture, 110μl PEG-calcium transfection solution (0.2M mannitol, 40% PEG4000, 100mM CaCl_2_) were added and mixed gently but completely. After incubation at room temperature for 5-15min, transfection process was terminated by adding 400-440μl W5 solution. The transfection mixture was then centrifuged at 100g for 2 min and the transfected protoplasts were collected, re-suspended in WI solution (4 mM MES, 0.5 M mannitol and 20 mM KCl, pH=5.7) with 1 mM DFHBI, examined and photographed using a Zeiss LSM 710 confocal laser scanning microscope (Yoo et al., 2007; Ding et al., 2009).

### *In Vitro* Transcription

Production of infectious PVX recombinant RNA transcripts was produced by in vitro transcription as described (Hong et al., 2001; Yu et al., 2020). Briefly, plasmid DNA of PVX/KK, PVX/KSK*1, PVX/KSK*2, PVX/KSK*3, PVX/KSK*4, PVX/KSK*5 and PVX/GFP was linearized by *Spe*I whilst pMD19-T/KK and pMD19-T/KSK plasmids were linearized by *PmlI*. The final concentration of purified linear plasmid DNA was 0.25 μg/μl. In vitro transcription was performed using 2.5 μg linear plasmid DNA as template and T7 RNA polymerase (NBL). Purified RNA transcripts were routinely dissolved in 40 μl in RNase-free water (Hong et al., 2001; Yu et al., 2020). Ten microliters of in vitro PVX RNA transcripts were mixed with 10-μl 200 μM DFHBI, incubated at 75°C for 5 min, then immediately cooled on ice, examined and photographed using a Nikon fluorescent stereomicroscope. The rest of RNA transcripts was used for virus-based RNA mimicking GFP assays in plants.

### Virus-based RMG Assay

Young leaves of *N. benthamiana* plants at six-leaf stage were inoculated with PVX recombinant RNA transcripts as described (Qin et al., 2017). In brief, the fourth and fifth fully expanded young leaves were dusted with a thin layer of carborundum (quartz sand, approximately 500 μm in diameter). A total of 20 μl viral RNA transcripts was dropped onto the two young leaves. Mechanical inoculation was then performed by gently finger-rubbing these leaves. Chlorotic lesions on inoculated leaves, characteristic of local PVX infection, typically appeared 3-4 days post inoculation (DPI), and systemic chlorosis developed on young leaves at 7 DPI and onwards. At 14-17 DPI, leaves showing clear chlorotic lesions were collected, cryo-sectioned, examined and photographed in the presence of 1mM DFHBI using a Zeiss LSM 710 confocal laser scanning microscope. At 17 DPI, total RNA was extracted from young leaves showing visible viral chlorosis symptoms using Qiagen RNeasy Plant Mini Kit. Ten milligrams of total RNA were mixed with 1 mM DFHBI, incubated at 75°C for 5 min, then immediately cooled on ice, examined and photographed using a Nikon fluorescent stereomicroscope equipped with a DN100 Digital Internet Camera (Nikon) (Paige et al., 2011).

### Quantitative RMG assay

i. Preparation of FITC solutions: To prepare 1 mg/ml (2.568×10^3^ μM) FITC stock solution, 1 mg fluorescein isothiocyanate isomer I powder (FITC, MW=389.38, Sigma-Aldrich) was dissolved in 1ml DMSO. We then prepared 5-ml 500 μM FITC solution by mixing 973.52 μl FITC stock solution with 4,026.48 μl Phosphate-Buffered Saline (PBS, 1.06mM KH_2_PO4, 155.17 mM NaCl, 2.97 mM Na_2_HPO_4_, pH= 7.4). Through a series of dilution in PBS by a factor of two, we prepared 5-ml FITC solution of 250 μM, 125 μM, 62.5 μM, 31.25 μM, 15.625 μM, 7.8125 μM, 3.90625 μM, 1.95 μM, 0.976 μM, 0.488 μM, 0.244 μM, and 0.122 μM, respectively. FITC fluorescence was observed under a Nikon fluorescent stereomicroscope (SMZ1500) and photographed using a DN100 Digital Internet Camera (Nikon) through a FITC (fluorescein isothiocyanate) filter or under transmitted white light (Supplemental Fig. S4A-C).
ii. FITC Standard Curve: For each concentration point, the intensity of FITC was measured in three – six biological duplicates using Fluorescence Spectrophotometer DS-11 FX (Denovix). PBS was used as a negative control. Fluorescent value was determined as a relative fluorescence unit (RFU) under long-pass emission at 514-567 nm and blue-led excitation at approximately 470nm. Oversaturation of fluorescence at higher concentration of the FITC solutions was clearly visible (Supplemental Fig. S4A and B). This phenomenon was reflected by abnormal RFU reading, likely due to out of detection limit of the spectrophotometer (Supplemental Fig. S5). Nevertheless, a standard curve could be drawn from our measurements from which an equation for RFU (Y-axis) vs FITC concentration (X-axis) was deduced (Supplemental Fig. S5).
iii. Quantitative RMG assay: A highly correlated equation y = 79297x – 9698 (R^2^ = 0.9978) was formulated by the Excel built-in software to quantitate RMG equivalent to the concentration range of 0 - 7.8125 μM FITC (Supplemental Fig. S5), where x represents the FITC micromolar concentration (μM) and y represents RFU. Based on the RFU readings for in vitro RNA transcripts or total RNA extracted from leaf tissues, we used this equation and calculated the strength of in vitro and ex vivo RMG in Fig. 5F; Fig. 6F and Supplemental Fig. S3.

### Nuclear acid Extraction, PCR and RT-PCR

Total RNA was extracted from leaf tissues using the RNeasy Plant Mini Kit (Qiagen). First-strand cDNA was synthesized using a Fast Quant RT Kit with gDNA Eraser (Tiangen). Genomic DNA was isolated from transgenic leaf tissues using the DNeasy Plant Mini Kit (Qiagen). RT-PCR or genomic PCR were performed using specific primers (Supplemental Table S1). *GAPDH* gene was used as an internal control (Qin et al., 2012).

## Data availability

Data supporting the findings of this work are available within the paper and its Supplemental Information files.

## Supplemental Data

The following supplemental materials are available.

**Supplemental Data Set S1.** Sequence information

**Supplemental Table S1.** Primers used in this study

**Supplemental Figure S1.** Spinach-based RMG in onion epidermal cells.

**Supplemental Figure S2.** Transient expression of Spinach in rice protoplasts

**Supplemental Figure S3.** Quantitative fluorescence of RNAs

**Supplemental Figure S4.** Fluorescence of FITC solutions

**Supplemental Figure S5.** Standard curve - FITC concentration vs fluorescence intensity

**Supplemental Figure S6.** DNA geminivirus-based RMG

**Supplemental Figure S7.** Infection of *N. benthamiana* by recombinant PVX

**Supplemental Figure S8.** Staining of RNA but not DNA by DFHBI.

## ACKNOWLEDGEMENTS

We thank Samine R Jaffery for providing us the pET28-c/Spinach2 plasmid and precious advices on RMG; Jiankang Zhu for the CRIPSR/Cas9 plasmid; Chun Wang, Lan Shen, Xixun Hu and Kejian Wang for their technical support and advice on preparation of rice protoplasts; Tien-Shin Yu for constructive advice on RMG; David Baulcombe for the original *Potato virus X*-based vector.

## LITERATURE CITED

Abudayyeh OO, Gootenberg JS, Essletzbichler P, Han S, Joung J, Belanto JJ, Verdine V, Cox DBT, Kellner MJ, Regev A, Lander ES, Voytas DF, Ting AY, Zhang F (2017). RNA targeting with CRISPR-Cas13. Nature 550: 280–284.

Autour A, Westhof E, Ryckelynck M (2016). iSpinach: a fluorogenic RNA aptamer optimized for in vitro applications. Nucleic Acids Res 44: 2491–2500.

Aw SS, Tang MX, Teo YN, Cohen SM (2016). A conformation-induced fluorescence method for microRNA detection. Nucleic Acids Res 44: e92.

Bai JY, Luo Y, Wang X, Li S, Luo M, Yin M, Zuo YL, Li GL, Yao JY, Yang H, Zhang MD, Wei W, Wang ML, Wang R, Fan CH, Zhao Y (2020). A protein-independent fluorescent RNA aptamer reporter system for plant genetic engineering. Nat Commun 11: 3847.

Banerjee AK, Chatterjee M, Yu Y, Suh SG, Miller WA, Hannapel DJ (2006). Dynamics of a mobile RNA of potato involved in a long-distance signaling pathway. Plant Cell 18: 3443–3457.

Bertrand E, Chartrand P, Schaefer M, Shenoy SM, Singer RH, Long RM (1998). Localization of ASH1 mRNA particles in living yeast. Mol Cell 2: 437–445.

Bruce G, Gu M, Shi N, Liu Y, Hong Y (2011) Influence of retinoblastoma-related gene silencing on the initiation of DNA replication by African cassava mosaic virus Rep in cells of mature leaves in *Nicotiana benthamiana* plants. Virol J 8: 561.

Buhtz A, Springer F, Chappell L, Baulcombe DC, Kehr J (2008). Identification and characterization of small RNAs from the phloem of Brassica napus. Plant J 53: 739–749.

Carrington JC, Kasschau KD, Mahajan SK, Schaad MC (1996). Cell-to-Cell and Long-Distance Transport of Viruses in Plants. Plant Cell 8: 1669–1681.

Chen S, Tao L, Zeng L, Vega-Sanchez ME, Umemura K, Wang GL (2006). A highly efficient transient protoplast system for analyzing defence gene expression and protein-protein interactions in rice. Mol Plant Pathol 7: 417–427.

Chen W, Zhang Q, Kong J, Hu F, Li B, Wu C, Qin C, Zhang P, Shi N, Hong Y (2015). MR VIGS: MicroRNA-Based Virus-Induced Gene Silencing in Plants. In: Mysore K., Senthil-Kumar M. (eds) Plant Gene Silencing. Methods in Molecular Biology, vol 1287. Humana Press, New York, NY. https://doi.org/10.1007/978-1-4939-2453-0_11

Chen W, Zhang X, Fan Y, et al (2018). A genetic Nnetwork for systemic RNA silencing in plants. Plant Physiol. 176: 2700–2719.

Chen X, Zhang D, Su N, Bao B, Xie X, Zuo F, Yang L, Wang H, Jiang L, Lin Q, Fang M, Li N, Hua X, Chen Z, Bao C, Xu J, Du W, Zhang L, Zhao Y, Zhu L, Loscalzo J, Yang Y (2019). Visualizing RNA dynamics in live cells with bright and stable fluorescent RNAs. Nat Biotechnol 37:1287–1293.

Deeken R, Ache P, Kajahn I, Klinkenberg J, Bringmann G, Hedrich R (2008). Identification of Arabidopsis thaliana phloem RNAs provides a search criterion for phloem-based transcripts hidden in complex datasets of microarray experiments. Plant J 55:746–759.

Ding WN, Yu ZM, Tong YL, Huang W, Chen HM, Wu P (2009). A transcription factor with a bHLH domain regulates root hair development in rice. Cell Res. 19: 1309–1311.

Dolgosheina EV, Jeng SC, Panchapakesan SS, Cojocaru R, Chen PS, Wilson PD, Hawkins N, Wiggins PA, Unrau PJ (2014). RNA mango aptamer-fluorophore: a bright, high-affinity complex for RNA labeling and tracking. ACS Chem Biol 9: 2412–2420.

Dong X, Van Wezel R, Stanley J, Hong Y (2003) Functional characterization of the nuclear localization signal for a suppressor of posttranscriptional gene silencing. Journal of virology 77: 7026–7033.

Ehrhardt DW, Frommer WB (2012). New technologies for 21st century plant science. Plant Cell 24: 374–394.

Ellison EE, Nagalakshmi U, Gamo ME, Huang PJ, Dinesh-Kumar S, Voytas DF (2020). Multiplexed Heritable Gene Editing Using RNA Viruses and Mobile Single Guide RNAs. Nat Plants 6: 620–624.

Feng Z, Zhang B, Ding W, Liu X, Yang DL, Wei P, Cao F, Zhu S, Zhang F, Mao Y, Zhu JK (2013). Efficient genome editing in plants using a CRISPR/Cas system. Cell Res 23: 1229–1232.

Filipovska A, Rackham O (2012). Modular recognition of nucleic acids by PUF, TALE and PPR proteins. Mol Biosyst 8: 699–708.

Filonov GS, Moon JD, Svensen N, Jaffrey SR (2014). Broccoli: rapid selection of an RNA mimic of green fluorescent protein by fluorescence-based selection and directed evolution. J Am Chem Soc 136: 16299–16308.

Guet D, Burns LT, Maji S, Boulanger J, Hersen P, Wente SR, Salamero J, Dargemont C (2015). Combining Spinach-tagged RNA and gene localization to image gene expression in live yeast. Nat Commun 6: 8882.

Haywood V, Yu TS, Huang NC, Lucas WJ (2005). Phloem long-distance trafficking of GIBBERELLIC ACID-INSENSITIVE RNA regulates leaf development. Plant J 42: 49–68.

Hong Y, Davies DL, Van Wezel R, Ellerker BE, Morton A, Barbara D (2001). Expression of the immunodominant membrane protein of chlorantie-aster yellows phytoplasma in *Nicotiana benthamiana* from a potato virus X-based vector. Acta horticulturae 550: 409–415.

Hong Y, Saunders K, Hartley MR, Stanley J (1996). Resistance to geminivirus infection by virus-induced expression of dianthin in transgenic plants. Virology 220: 119–127.

Huang H, Suslov NB, Li NS, Shelke SA, Evans ME, Koldobskaya Y, Rice PA, Piccirilli JA (2014). A G-quadruplex-containing RNA activates fluorescence in a GFP-like fluorophore. Nat Chem Biol 10: 686–691.

Huang K, Doyle F, Wurz ZE, Tenenbaum SA, Hammond RK, Caplan JL, Meyers BC (2017). FASTmiR: an RNA-based sensor for in vitro quantification and live-cell localization of small RNAs. Nucleic Acids Res 45: e130.

Huang NC, Jane WN, Chen J, Yu TS (2012). Arabidopsis thaliana CENTRORADIALIS homologue (ATC) acts systemically to inhibit floral initiation in Arabidopsis. Plant J 72: 175–184.

Jackson SD, Hong Y (2012). Systemic movement of FT mRNA and a possible role in floral induction. Front Plant Sci 3: 127.

Kim G, LeBlanc ML, Wafula EK, dePamphilis CW, Westwood JH (2014). Genomic-scale exchange of mRNA between a parasitic plant and its hosts. Science 345: 808–811.

Lai T, Wang X, Ye B, Jin M, Chen W, Wang Y, Zhou Y, Blanks AM, Gu M, Zhang P, Zhang X, Li C, Wang H, Liu Y, Gallusci P, Tör M, Hong Y (2020) Molecular and functional characterization of the SBP-box transcription factor SPL-CNR in tomato fruit ripening and cell death. J Exp Bot 71: 2995–3011.

Li C, Gu M, Shi N, Zhang H, Yang X, Osman T, Liu Y, Wang H, Vatish M, Jackson S, Hong Y (2011). Mobile FT mRNA contributes to the systemic florigen signalling in floral induction. Sci Rep 1:73.

Li C, Zhang K, Zeng X, Jackson S, Zhou Y, Hong Y (2009). A cis element within flowering locus T mRNA determines its mobility and facilitates trafficking of heterologous viral RNA. J Virol 83: 3540–3548.

Liu L, Chen X (2018). Intercellular and systemic trafficking of RNAs in plants. Nat Plants 4: 869–878.

Lu KJ, Huang NC, Liu YS, Lu CA, Yu TS (2012). Long-distance movement of Arabidopsis FLOWERING LOCUS T RNA participates in systemic floral regulation. RNA Biol 9: 653–662.

Luo KR, Huang NC, Yu TS (2018). Selective targeting of mobile mRNAs to plasmodesmata for cell-to-cell movement. Plant Physiol 177: 604–614.

Ma X, Zhang Q, Zhu Q, Liu W, Chen Y, Qiu R, Wang B, Yang Z, Li H, Lin Y, Xie Y, Shen R, Chen S, Wang Z, Chen Y, Guo J, Chen L, Zhao X, Dong Z, Liu YG (2015). A Robust CRISPR/Cas9 System for Convenient, High-Efficiency Multiplex Genome Editing in Monocot and Dicot Plants. Mol Plant 8: 1274–1284.

Nelles DA, Fang MY, O’Connell MR, Xu JL, Markmiller SJ, Doudna JA, Yeo GW (2016). Programmable RNA Tracking in Live Cells with CRISPR/Cas9. Cell 165: 488–496.

Notaguchi M, Higashiyama T, Suzuki T (2015). Identification of mRNAs that move over long distances using an RNA-Seq analysis of Arabidopsis/Nicotiana benthamiana heterografts. Plant Cell Physiol 56: 311–321.

Paige JS, Wu KY, Jaffrey SR (2011). RNA mimics of green fluorescent protein. Science 333: 642–646.

Pothoulakis G, Ceroni F, Reeve B, Ellis T (2014). The spinach RNA aptamer as a characterization tool for synthetic biology. ACS Synth Biol 3: 182–187.

Qin C, Chen W, Shen J, Cheng L, Akande F, Zhang K, Yuan C, Li C, Zhang P, Shi N, Cheng Q, Liu Y, Jackson S, Hong Y (2017) A virus-induced assay for functional dissection and analysis of monocot and dicot flowering time genes. Plant Physiol. 174: 875–885.

Qin C, Shi N, Gu M et al (2012) Involvement of RDR6 in short-range intercellular RNA silencing in *Nicotiana benthamiana*. Sci Rep 2: 467.

Qin C, Zhang Q, He M, Kong J, Li B, Mohamed A, Chen W, Zhang P, Zhang X, Yu Z, et al. (2015) Virus technology for functional genomics in plants. In P Poltronieri and Y Hong, eds, Applied Plant Genomics and Biotechnology. Elsevier Woodhead Publishing, Cambridge, UK, pp 229–236.

Ruiz-Medrano R, Xoconostle-Cázares B, Lucas WJ (1999). Phloem long-distance transport of CmNACP mRNA: implications for supracellular regulation in plants. Development 126: 4405–4419.

Sainsbury F, Lomonossoff GP (2008). Extremely high-level and rapid transient protein production in plants without the use of viral replication. Plant Physiol 148: 1212–1218.

Sainsbury F, Thuenemann EC, Lomonossoff GP (2009). pEAQ: versatile expression vectors for easy and quick transient expression of heterologous proteins in plants. Plant Biotechnol J 7: 682–693.

Scheiba RM, de Opakua AI, Díaz-Quintana A, Cruz-Gallardo I, Martínez-Cruz LA, Martínez-Chantar ML, Blanco FJ, Díaz-Moreno I (2014). The C-terminal RNA binding motif of HuR is a multi-functional domain leading to HuR oligomerization and binding to U-rich RNA targets. RNA Biol 11: 1250–1261.

Schönberger J, Hammes UZ, Dresselhaus T (2012). In vivo visualization of RNA in plants cells using the λN22 system and a GATEWAY-compatible vector series for candidate RNAs. Plant J 71:173–181.

Shahid S, Kim G, Johnson NR, Wafula E, Wang F, Coruh C, Bernal-Galeano V, Phifer T, dePamphilis CW, Westwood JH, Axtell MJ (2018). MicroRNAs from the parasitic plant Cuscuta campestris target host messenger RNAs. Nature 553: 82–85.

Shi Y, Gu M, Fan Z, Hong Y (2008) RNA silencing suppressors: how viruses fight back. Future Virol 3: 125–133.

Strack RL, Disney MD, Jaffrey SR (2013). A superfolding Spinach2 reveals the dynamic nature of trinucleotide repeat-containing RNA. Nat Methods 10: 1219–1224.

Tang Y, Wang F, Zhao J, Xie K, Hong Y, Liu Y (2010). Virus-based microRNA expression for gene functional analysis in plants. Plant Physiol 153: 632–641.

Thieme CJ, Rojas-Triana M, Stecyk E, Schudoma C, Zhang W, Yang L, Miñambres M, Walther D, Schulze WX, Paz-Ares J, Scheible WR, Kragler F (2015). Endogenous Arabidopsis messenger RNAs transported to distant tissues. Nat Plants 1: 15025.

Tutucci E, Livingston NM, Singer RH, Wu B (2018). Imaging mRNA In Vivo, from Birth to Death. Annu Rev Biophys 47: 85–106.

Tyagi S, Kramer FR (1996). Molecular beacons: probes that fluoresce upon hybridization. Nat Biotechnol 14: 303–308.

Uddin MN, Kim JY (2013). Intercellular and Systemic Spread of RNA and RNAi in Plants. Wiley Interdiscip Rev RNA 4: 279–293.

van Wezel R, Liu H, Tien P, Stanley J, Hong Y (2001) Gene C2 of the monopartite geminivirus tomato yellow leaf curl virus-China encodes a pathogenicity determinant that is localized in the nucleus. Mol Plant Microbe Interact 14: 1125–1128.

Wang X, McLachlan J, Zamore PD, Hall TM (2002). Modular recognition of RNA by a human pumilio-homology domain. Cell 110: 501–512.

Warner KD, Chen MC, Song W, Strack RL, Thorn A, Jaffrey SR, Ferré-D’Amaré AR (2014). Structural basis for activity of highly efficient RNA mimics of green fluorescent protein. Nat Struct Mol Biol 21:658–663.

Warner KD, Sjekloća L, Song W, Filonov GS, Jaffrey SR, Ferré-D’Amaré AR (2017). A homodimer interface without base pairs in an RNA mimic of red fluorescent protein. Nat Chem Biol 13: 1195–1201.

Wydro M, Kozubek E, Lehmann P (2006). Optimization of transient Agrobacterium-mediated gene expression system in leaves of *Nicotiana benthamiana*. Acta Biochim Pol 53: 289–298.

Xoconostle-Cázares B, Xiang Y, Ruiz-Medrano R, Wang HL, Monzer J, Yoo BC, McFarland KC, Franceschi VR, Lucas WJ (1999). Plant paralog to viral movement protein that potentiates transport of mRNA into the phloem. Science 283: 94–98.

Yamada T, Yoshimura H, Inaguma A, Ozawa T (2011). Visualization of nonengineered single mRNAs in living cells using genetically encoded fluorescent probes. Anal Chem 83: 5708–5714.

Yang HW, Yu TS (2010). Arabidopsis floral regulators FVE and AGL24 are phloem-mobile RNAs. Botanical Studies 51: 17–26.

Yoo BC, Kragler F, Varkonyi-Gasic E, Haywood V, Archer-Evans S, Lee YM, Lough TJ, Lucas WJ (2004). A systemic small RNA signaling system in plants. Plant Cell 16: 1979–2000.

Yoo SD, Cho YH, Sheen J (2007). Arabidopsis mesophyll protoplasts: a versatile cell system for transient gene expression analysis. Nat Protoc 2: 1565–1572.

Yoshida S, Forno DA, Cock JH, Gomez KA (1976). Laboratory manual for physiological studies of rice, 3rd edn. Manila, The Philippines: International Rice Research Institute.

Yoshimura H, Inaguma A, Yamada T, Ozawa T (2012). Fluorescent probes for imaging endogenous beta-actin mRNA in living cells using fluorescent protein-tagged pumilio. ACS Chem Biol 7: 999–1005.

You M, Jaffrey SR (2015). Structure and Mechanism of RNA Mimics of Green Fluorescent Protein. Annu Rev Biophys 44:187–206.

Yu Z, Cho SK, Zhang P, Hong Y, Hannapel DJ (2020). Utilizing Potato Virus X to Monitor RNA Movement. RNA Tagging. Methods in Molecular Biology, pp:181–194.

Yu Z, Dong L, Jiang Z, Yi K, Zhang J, Zhang Z, Zhu Z, Wu Y, Xu M, Ni J (2018). A semi-dominant mutation in a CC-NB-LRR-type protein leads to a short-root phenotype in rice. Rice 11: 54.

Zhang J, Fei J, Leslie BJ, Han KY, Kuhlman TE, Ha T (2015) Tandem Spinach Array for mRNA Imaging in Living Bacterial Cells. Sci Rep 5: 17295.

Zhang X, Lai T, Zhang P, Zhang X, Yuan C, Jin Z, Li H, Yu Z, Qin C, Tör M, Ma P, Cheng Q, Hong Y (2019) Mini review: Revisiting mobile RNA silencing in plants. Plant Sci. 278: 113–117.

