## Supplemental Data for "Reappraisal of Spinach-based RNA Visualization in Plants"

[1] **Funding:** This work was in part supported by grants from Ministry of Agriculture of the People's Republic of China (National Transgenic Program of China 2016ZX08009001-004); National Natural Science Foundation of China (31200913, 31872636); Zhejiang Provincial Natural Science Foundation (LY19C020002), China Scholarship Council (201709645003), the Entrepreneurship and Innovation Project for the Overseas Returnees (or Teams) in Hangzhou (4105C5062000611). Ministry of Science & Technology of China (National Key R&D Program 2017YFE0110900); Hangzhou Normal University (Sino-EU Plant RNA Signaling S&T Platform Initiative 9995C5021841101).

The author responsible for distribution of materials integral to the findings presented in this article in accordance with the policy described in the instructions for Authors (www.plantphysioll.org) is: Yiguo Hong (, and).

**Author Contributions:** Z.Y. designed, performed all experiments, analyzed data and drafted the manuscript; F.M. H.Y. and Q.C. performed plant transformation, plant RNA extraction and RT-PCR analysis. M.Y., S.L., Y.W., X.Z., and P.Z. performed virus-based RMG assays; S.J., N.S., and Y.L. were involved in the analysis of data and helped write the article; Y.H. initiated the project, conceived experiments, analyzed data, and wrote the article.

**Competing interests:** The authors declare no competing interests.

**One sentence summary:** Spinach-based RMG technology was re-evaluated to have potential for ex vivo and in vivo monitoring RNAs in plant cells.

### Supplemental Data Set

**Supplemental Data Set S1.** Sequence information

The original sequences used for all constructs (Fig. 1A; Fig. 2A; Fig. 4A; Fig. 5A; Supplemental Fig. 1A and Supplemental Fig. 5A) are shown below.

(i) *AttRNA*$^{Lys}$ (abbreviated to K*,* 73 nt):

5’-GCCCGTCTAG-CTCAGTTGGT-AGAGCGCAAG-GCTCTTAACC-TTGTGGTCGT-GGGTTCGAGC-CCCACGGTGG-GCG-3’

(ii) *Spinach* sequence (abbreviated to S, 80 nt):

5’-GACGCGACCG-AAATGGTGAA-GGACGGGTCC-AGTGCTTCGG-CACTGTTGAG-TAGAGTGTGA-GCTCCGTAAC-TGGTCGCGTC-3’

(iii) *AttRNA*$^{Lys}$-*AttRNA*$^{Lys}$ (abbreviated to KK) sequence (152 nt):

5’-GCCCGTCTAG-CTCAGTTGGT-AGAGCGCAAG-GCTCTTAACC-TTGTGGTCGT-GGGTTCGAGC-CCCACGGTGG-GCG**cggccg**G-CCCGTCTAGC-TCAGTTGGTA-GAGCGCAAGG-CTCTTAACCT-TGTGGTCGTG-GGTTCGAGCC-CCACGGTGGG-CG-3’ (The underline sequence is the *Eag* I site.)

(iv) *AttRNA*$^{Lys}$-*Spinach*-*AttRNA*$^{Lys}$ (abbreviated to KSK) sequence (250 nt):

5’-accggt**GCCC-GTCTAGCTCA-GTTGGTAGAG-CGCAAGGCTC-TTAACCTTGT-GGTCGTGGGT-TCGAGCCCCA-CGGTGGGCG**a agctt**GACGC-GACCGAAATG-GTGAAGGACG-GGTCCAGTGC-TTCGGCACTG-TTGAGTAGAG-TGTGAGCTCC-GTAACTGGTC-GCGTC**gcatg-c**GCCCGTCTA-GCTCAGTTGG-TAGAGCGCAA-GGCTCTTAAC-CTTGTGGTCG-TGGGTTCGAG-CCCCACGGTG-GGCG**cccggg-3’ (The underline sequence from 5’ to 3’ is *Age* I, *Hind* III, *Sph* I and *Sma* I (*Xma* I) site, respectively. The green sequence is *Spinach1*. The bold sequence on each side of *Spinach* is *AttRNA*$^{Lys}$.)

(v) T7/KK sequence (227 nt):

5’-taatacgactcactatagggTCACCACCAC-GGAATCGATacgcgt**GCCCG-TCTAGCTCAG-TTGGTAGAGC-GCAAGGCTCT-TAACCTTGTG-GTCGTGGGTT-CGAGCCCCAC-GGTGGGCG**cggccg**GCCCGT-CTAGCTCAGT-TGGTAGAGCG-CAAGGCTCTT-AACCTTGTGG-TCGTGGGTTC-GAGCCCCACG-GTGGGCGTCC-GGATGATATC-GTCGACCGCC-G**cacgtg-3’ (The sequence in the box is the T7 promoter. The underline sequence from 5’ to 3’ is *Mlu* I, *Eag* I, and *Pml* I site respectively.)

(vi) T7/K-Spinach-K sequence (KSK, 374 nt)

5’-taatacgactcactatagggTCACCACCAC-GGAATCGATacgcgtATATT-CTGCCCAAAT-TCGCGaccggt**GCCCGTCTA-GCTCAGTTGG-TAGAGCGCAA-GGCTCTTAAC-CTTGTGGTCG-TGGGTTCGAG-CCCCACGGTG-GGCG**aagctt**GACGCGACCG-AAATGGTGAA-GGACGGGTCC-AGTGCTTCGG-CACTGTTGAG-TAGAGTGTGA-GCTCCGTAAC-TGGTCGCGTC-**gcatgc**GCCC-GTCTAGCTCA-GTTGGTAGAG-CGCAAGGCTC-TTAACCTTGT-GGTCGTGGGT-TCGAGCCCCA-CGGTGGGCG**cccgggCATCA-CCATCACCAT-CACTACGGCC-GTGATCCGGA-TGATATCGTC-GACCGCCGca cgtg-3’ (The sequence in the box is the T7 promoter. The underline sequence from 5’ to 3’ is *Mlu* I, Age I, *Hind* III, *Sph* I, *Sma* I (*Xma* I) and *Pml* I site respectively. The green sequence is Spinach. The bolded sequence on each side of *Spinach* is *AttRNA*$^{Lys}$.)

(vii) pCVA/KK sequence (205 nt):

5’-ggtaccACCA-CGGAATCGAT-acgcgt**GCCC-GTCTAGCTCA-GTTGGTAGAG-CGCAAGGCTC-TTAACCTTGT-GGTCGTGGGT-TCGAGCCCCA-CGGTGGGCG**cggccg**GCCCG-TCTAGCTCAG-TTGGTAGAGC-GCAAGGCTCT-TAACCTTGTG-GTCGTGGGTT-CGAGCCCCAC-GGTGGGCG**tc-cggaTGATAT-CGTCGACCGtctaga-3’ (The underline sequence from 5’ to 3’ is *Kpn* I, *Mlu* I, *Eag* I, *BspE* I and *Xba* I site, respectively. The bold sequences are *AttRNA*$^{Lys}$)

(viii) pCVA/KSK sequence (352 nt)

5’-ggtaccACCA-CGGAATCGAT-ACGCGTATAT-TCTGCCCAAA-TTCGCGaccggt**GCCCGTCT-AGCTCAGTTG-GTAGAGCGCA-AGGCTCTTAA-CCTTGTGGTC-GTGGGTTCGA-GCCCCACGGT-GGGCG**aagct-t**GACGCGACC-GAAATGGTGA-AGGACGGGTC-CAGTGCTTCG-GCACTGTTGA-GTAGAGTGTG-AGCTCCGTAA-CTGGTCGCGT-C**gcatgc**GCC-CGTCTAGCTC-AGTTGGTAGA-GCGCAAGGCT-CTTAACCTTG-TGGTCGTGGG-TTCGAGCCCC-ACGGTGGGCG-**cccgggCATC-ACCATCACCA-TCACTACGGC-CGTGATCCGG-ATGATATCGT-CGACCGtctaga-3’

(The underline sequence from 5’ to 3’ is *Kpn* I, *Age* I, *Hind* III, *Sph* I, Sma I/ (Xma I) and *Xba* I site, respectively. The green sequence is *Spinach.* The bold sequence on each side of *Spinach* is *AttRNA*$^{Lys}$).

### Supplemental Figures

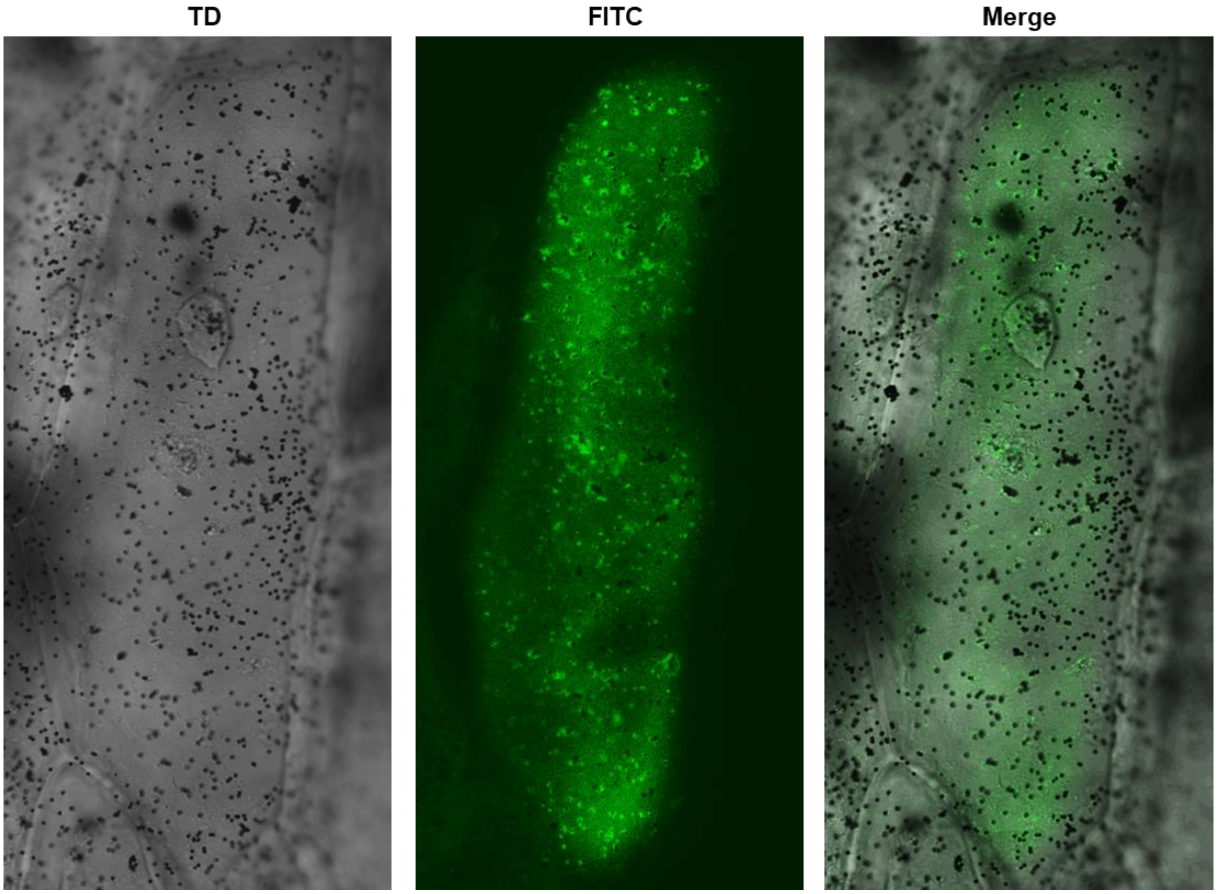


**Supplemental Figure S1.** Spinach-based RMG in onion epidermal cells. The onion cell in Fig. 2F was enlarged to show RMG in detail. Under transmitted white light (TD), numerous gold particles coated with pEAQ-HT/KSK was visible as “dark” dots (Left panel). Strong green fluorescence was observed under the FITC filter (middle panel). In the merged image (right) clearly shows green fluorescence around the “dark” gold particle and throughout the cytoplasm of this onion epidermal cell. Photographed was taken as described in the legend of Fig. 2.

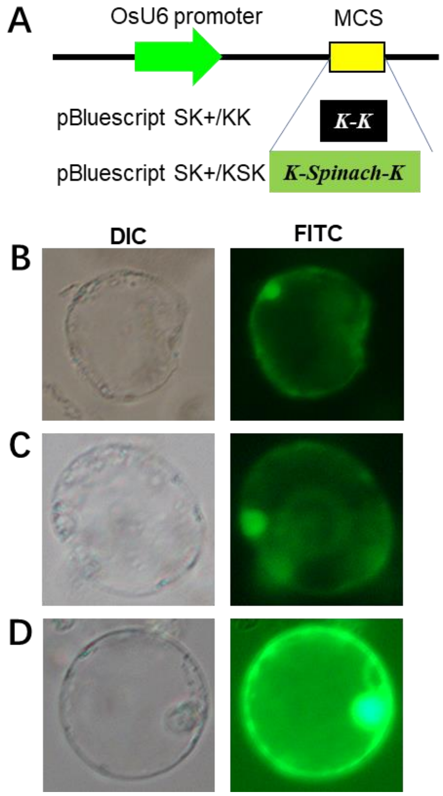


**Supplemental Figure S2.** Transient expression of Spinach in rice protoplasts. A, Diagrammatic of OsU6:KK and OsU6:KSK expression cassette in pBluescript SK+/OsU6 vector. Expression of KK and KSK is under the control of the rice U6 promoter. B-C, RMG in rice protoplasts. Protoplasts were transfected with pBluescript SK+/KK (B), pBluescript SK+/KSK (C) or pBluescript SK+/GFP (D) as positive control. Photographs were taken through DIC (Right panel) or FITC filter (Left panel).

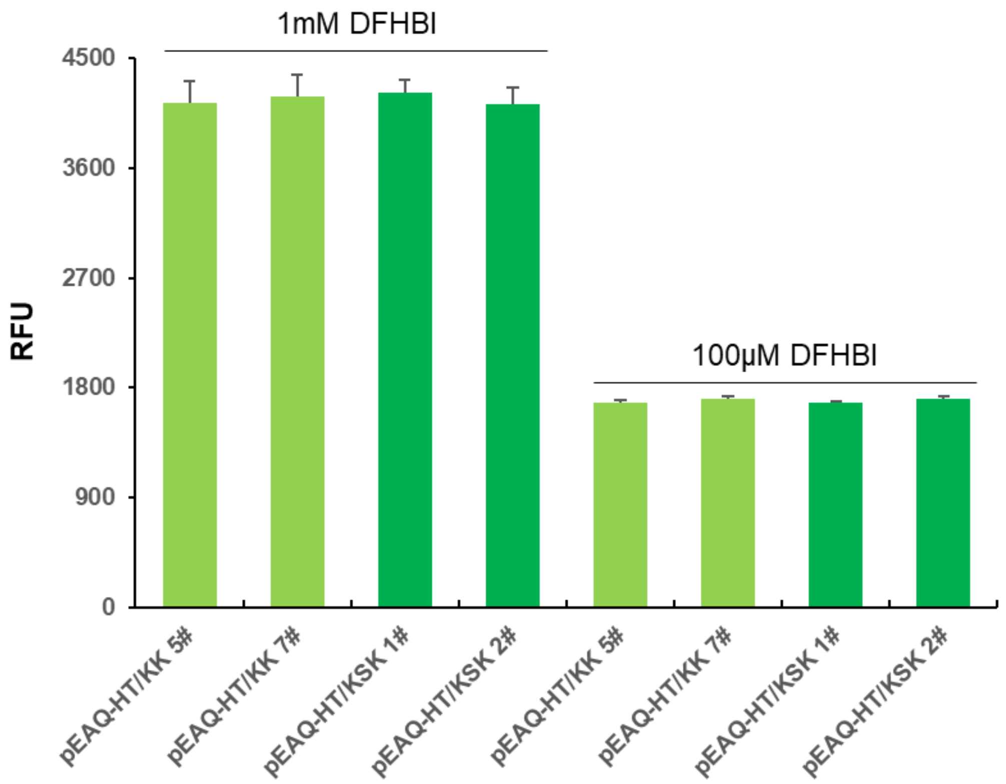


**Supplemental Figure S3**. Quantitative fluorescence of RNAs. Total RNA was extracted from transgenic KK lines 5# and 7#, and KSK lines 1# and 2#. RNA fluorescence was measured as relative fluorescence units (RFU) at the 1 mM or 100 µM DFHBI. Using the equation based on the FITC vs fluorescence standard curve (Supplemental Fig. S3A-C and Supplemental Fig. S4), we calculated the average concentration equivalent to FITC was $1.39 \times 10^{-4}$ µM for total RNAs extracted from the *KSK* transgenic leaf tissues after deducting the background fluorescence in transgenic *KK* plant RNA samples.

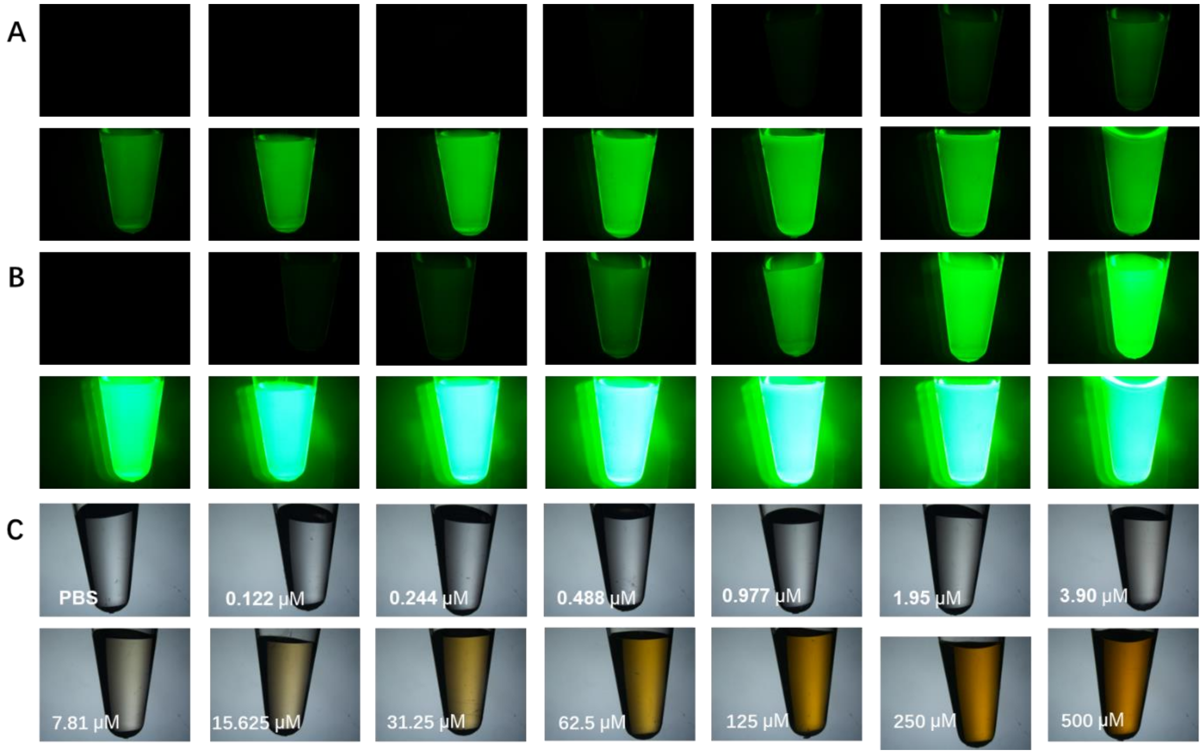


**Supplemental Figure S4.** Fluorescence of FITC solutions. A and B, Fluorescent exposure time was 400 milliseconds (A), or 6 seconds (B). C, Photographic record of FITC samples through transmitted detector. FITC concentration (µM) in PBS for each solution is shown C. The same order for each of the FITC solution was arranged in A and B. PBS was used as negative control.

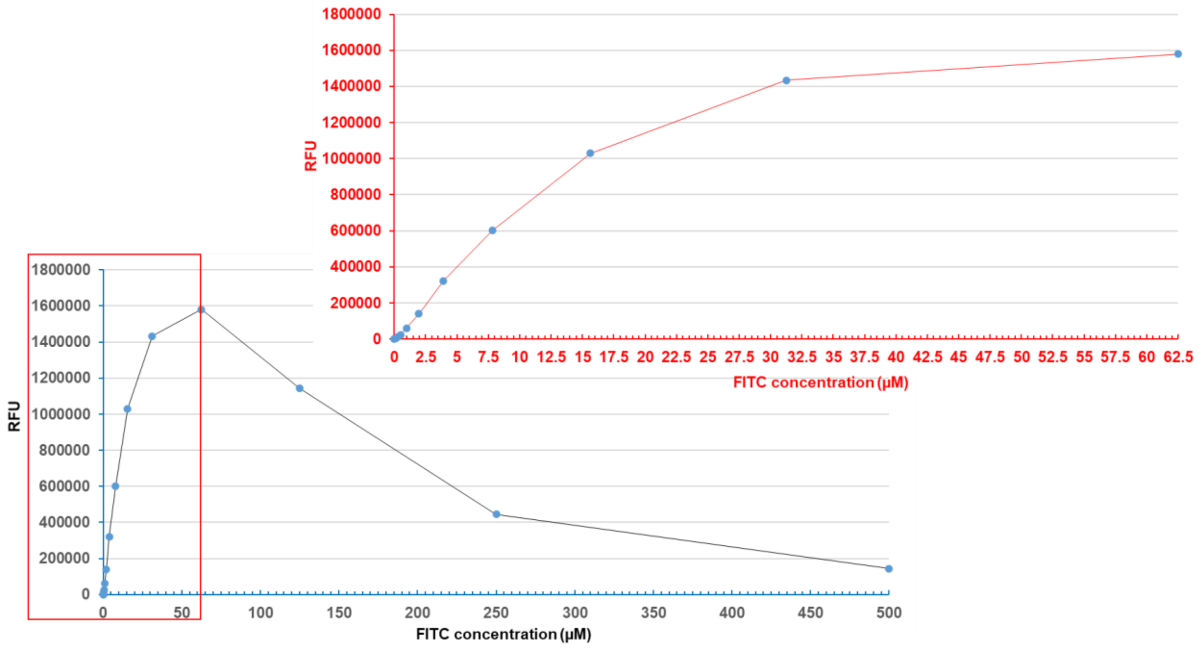


**Supplemental Figure S5**. Standard curve for FITC concentration vs fluorescence intensity. A plot of FITC concentration vs fluorescence. X-axis shows the concentration of FITC ranging from 0 to 500 µM. Y-axis is fluorescence value of relative fluorescence units (RFU, 514-567nm) measured by a fluorescence spectrophotometer (Denovix). The boxed section of the curve is enlarged to show a linear correlation between fluorescent intensity and FITC concentration ranging from 0 to 62.5 µM. A highly correlated equation y = 79297x – 9698 ($R^2$ = 0.9978) was formulated by the Excel built-in software to quantitate RMG equivalent to the concentration range of 0 - 7.8125 µM FITC. For each concentration point, 3 – 6 biological duplicates were used to measure RFU (Mean ± SD). Mean value was used to plot the FITC standard curve.

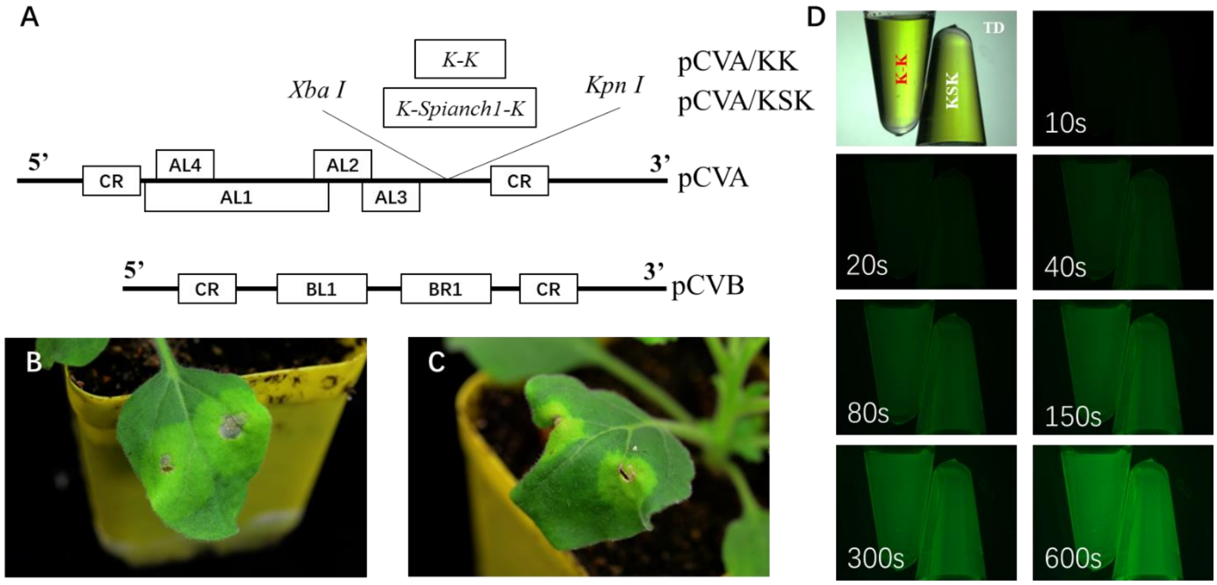


**Supplemental Figure S6**. DNA geminivirus-based RMG. A, Diagrammatic of bipartite cabbage leaf curl virus DNA A-based vector pCVA (Tang et al., 2010; Chen et al., 2015). The genome of viral DNA B (pCVB) is indicated. KK and KSK were cloned into the *Xba*I/*Kpn*I sites and replaced the viral coat protein gene in pCVA. Viral genomic DNA A encodes 4 complementary genes, namely *AL1* (Replication-associated protein), *AL2* (Transcriptional activator), *AL3* (Replication-enhancer) and *AL4*; while DNA-B encodes 2 genes BR1 and BL1 for nuclear-shuttle and movement proteins. CR stands for common region which consists of almost identical nucleotide sequences between viral DNA A and DNA B. It consists of viral DNA replication origin as well as bi-directional promoters for controlling expression of genes located on the viral sense and complementary strands of DNA-A and DNA-B. B and C, Agroinfiltration assays. Young leaves of *N. benthamiana* plants were infiltrated with a mixture of agrobacterium harboring pCVB and pCVA/KK (B) or pCVA/KSK (C). Photographs were taken at 8-days post-agroinfiltration (DPA). D, *Ex planta* RMG assay. Five micrograms of total RNA extracted from agroinfiltrated leaves at 8 DPA in 2 mM DFHBI were photographically recorded under transmitted light (TD) or FITC filter. Exposure time ranged from 10 to 600 seconds (s) as indicated in each image. The positions of KK and KSK samples in all images are same as indicated in the TD-labeled one (D).

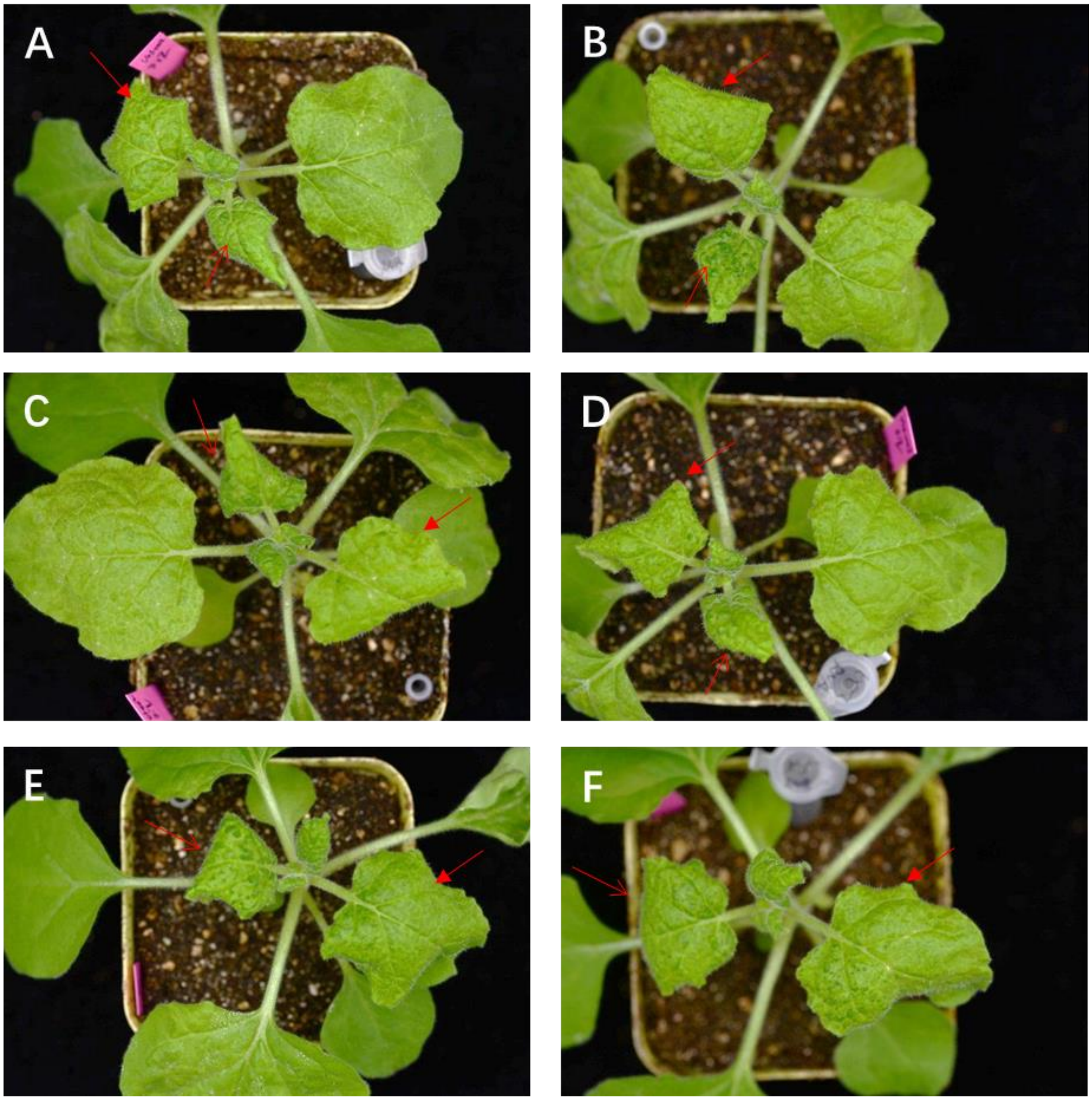


**Supplemental Figure S7.** Infection of *N. benthamiana* by recombinant PVX. A-F, Development of viral systemic symptoms. *N. benthamiana* plants inoculated with *in vitro* RNA transcripts for PVX/KK (A), PVX/KSK*1 (B), PVX/KSK*2 (C), PVX/KSK*3 (D), PVX/KSK*4 (E), or PVX/KSK*5 (F) developed chlorosis on newly grown young leaves (red arrow). Plants were photographed at 17 days post-inoculation.

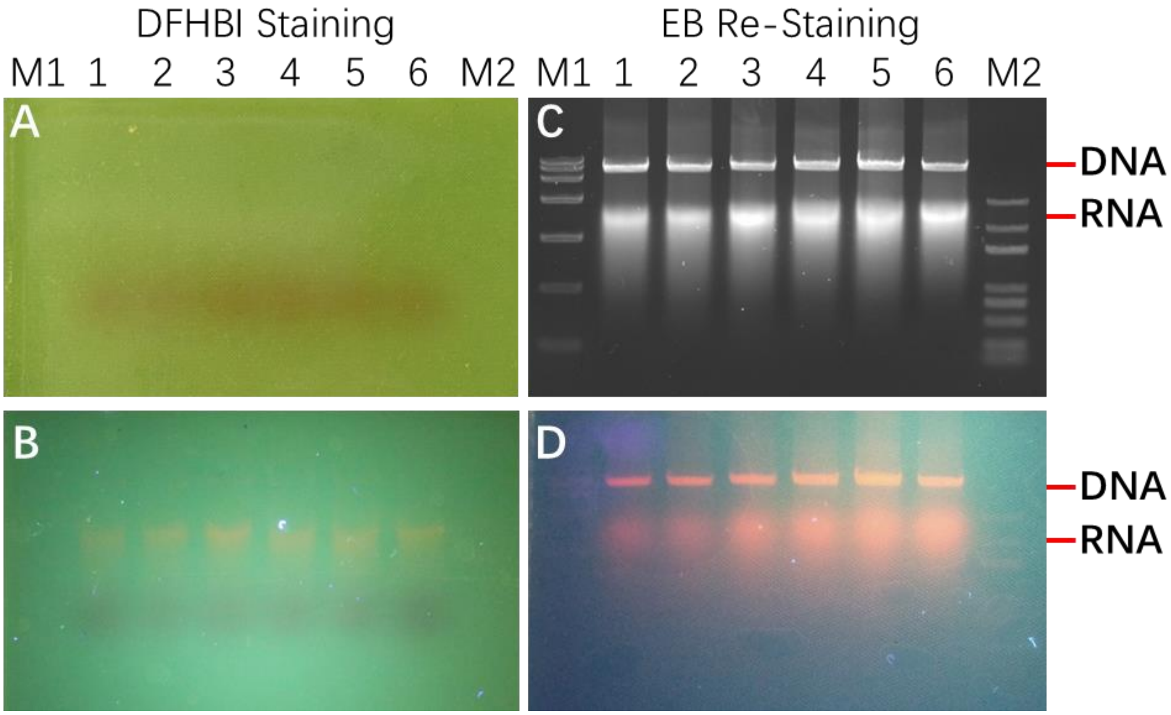


**Supplemental Figure S8.** Specific staining of RNA by DFHBI. A, DFHBI-staining gel under visible light. Samples loaded on this gel were same as these in Fig. 4B. B, DFHBI-staining gel under long-wavelength ultraviolet (UV) light. C, Ethidium Bromide (EB) re-staining gel under short-wavelength UV (also same as Fig. 4B). D, EB re-staining gel under long-wavelength UV. Images A, B and D were taken with a Canon EOS R Mirrorless Digital Camera (24-105mm f/4L Lens) through a UV filter. Image C was taken using the Bio-Rad gel imaging analysis system. M1 and M2: BM15000 and BM5000 DNA markers (Biomed); Lanes 1 and 2: PVX/KK; Lane 3: PVX/KSK; Lane 4: PVX/AtFT:KSK; Lane 5: PVX/mAtFT:KSK; Lane 6: PVX/AtTFL1:KSK. After terminating *in vitro* transcription (IVT) reactions, 5 µl of IVT mixtures were run on 1% Agarose-TAE gel. Gel was then immersed in 200 µM DFHBI solution for 15 minutes and photographically recorded. Then the same gel was rinsed three times in RNase-free water for 30 minutes, re-stained in 10 µg/ml ethidium bromide (EB) solution and photographed. We found that DFHBI could only specifically stain RNA transcripts tagged with or without spinach (B) although EB can stain both RNA and DNA (C and D).

**Supplemental Table S1. Primers used in this study**

| Primer code | Primer name | Primer sequence (underline sequences are Restriction Endonuclease sites) | Use |
|---|---|---|---|
| P001 | PVX/K-K (1) For | 5'-**CGCGT**GCCCGTCTAGCTCAGTTGGTAGAGCGCAAGGCTCTTAACCTTGTGGTCGTGGGTTCGAGCCCCACGGTGGGCG**C**-3' | KK template (4-1) for PVX/KK construction containing partial Mlu I and Eag I sites nucleotides |
| P002 | PVX/K-K (2) Rev | 5'-**GGCCG**CGGGCAGATCGAGTCAACCATCTCGCGTTCCGAGAATTGGAACACCAGCACCCAAGCTCGGGGTGCCACCCGC**A**-3' | KK template (4-2) for PVX/KK construction containing partial Mlu I and Eag I sites nucleotides |
| P003 | PVX/K-K (3) For | 5'-**GGCCG**GCCCGTCTAGCTCAGTTGGTAGAGCGCAAGGCTCTTAACCTTGTGGTCGTGGGTTCGAGCCCCACGGTGGGCG**T**-3' | KK template (4-3) for PVX/KK construction containing partial Mlu I and Eag I sites nucleotides |
| P004 | PVX/K-K (4) Rev | 5'-**CCGGA**CGGGCAGATCGAGTCAACCATCTCGCGTTCCGAGAATTGGAACACCAGCACCCAAGCTCGGGGTGCCACCCGC**C**-3' | KK template (4-4) for PVX/KK construction containing partial Mlu I and Eag I sites nucleotides |
| P005 | pEAQ-HT/K-K For | 5'-ATA**TCGCGA**CAATCACAGTGTTGGCTTGCAAAC-3' | For pEAQ-HT/KK construction with Nru I site |
| P006 | pEAQ-HT/K-K Rev | 5'-ATA**CTCGAG**TTGACCCTATGGGCTGTGTTG-3' | For pEAQ-HT/KK construction with Xho I site |
| P007 | pMD19-T/T7:K-Spinach1-K (T7:K-K) For | 5'-**TAATACGACTCACTATAGGG**TCACCACCACGGAATCGATACGC-3' | For pMD19-T/KK (or KSK) construction with T7 promoter sequence |
| P008 | pMD19-T/T7:K-Spinach1-K (T7:K-K) Rev | 5'-ATA**CACGTG**CGGCGGTCGACGATATCATCCG-3' | For pMD19-T/KK or KSK construction with Pml I site |
| P009 | PVX/K-Spinach1-K(1) For | 5'-ATA**ACGCGT**ATATTCTGCCCAAATTCGCGA | For PVX/K-Spinach1-K(1) construction with Mlu I site |
| P010 | PVX/K-Spinach1-K(1) Rev | 5'-ATA**CGGCCG**TAGTGATGGTGATGGTGATGCCCGGG | For PVX/K-Spinach1-K(1) construction with Eag I site |
| P011 | PVX/K-Spinach1-K(2) For | 5'-ATA**CGGCCG**ATATTCTGCCCAAATTCGCGA | For PVX/K-Spinach1-K(2) construction with Eag I site |
| P012 | PVX/K-Spinach1-K(2) Rev | 5'-ATA**GTCGAC**TAGTGATGGTGATGGTGATGCCCGGG | For PVX/K-Spinach1-K(2) construction with Sal I site |
| P013 | PVX/AtFT-K-Spinach1-K For | 5'-ATA**ATCGAT**ATGTCTATAAAATATAAGAGACCCTCTTATAGT-3' | For PVX/AtFT-KSK construction with Cla I site |
| P014 | PVX/AtFT-K-Spinach1-K Rev | 5'-ATA**CGGCCG**CTAAAGTCTTCTTCCTCCGCAG-3' | For PVX/AtFT-KSK construction with Eag I site |
| P015 | PVX/AtmFT-K-Spinach1-K For | 5'-ATA**ATCGAT**TAGTCTATAAAATATAAGAGACCCTCTTATAGT-3' | For PVX/AtmFT-KSK construction with Cla I site |
| P016 | PVX/AtmFT-K-Spinach1-K Rev | 5'-ATA**CGGCCG**CTAAAGTCTTCTTCCTCCGCAG-3' | For PVX/AtmFT-KSK construction with Eag I site |
| P017 | PVX/AtTFL1-K-Spinach1-K For | 5'-ATA**ATCGAT**ATGGAGAATATGGGAACTAGAGTGATAGAGC-3' | For PVX/AtTFL1-KSK construction with Cla I site |
| P018 | PVX/AtTFL1-K-Spinach1-K Rev | 5'-ATA**ACGCGT**CTAGCGTTTGCGTGCAGCGGT-3' | For PVX/AtTFL1-KSK construction with Mlu I site |
| P019 | pCVA/K-Spinach1-K (K-K) For | 5'-ATA**GGTACC**ACCACGGAATCGATACGCGT-3' | For pCVA/KSK (or KK) construction with Kpn I site |
| P020 | pCVA/K-Spinach1-K(K-K) Rev | 5'-ATA**TCTAGA**CGGTCGACGATATCATCCGGAT-3' | For pCVA/KSK (or KK) construction with Xba I site |
| P021 | pMD19-T/K-Spinach1-K (Cla I) For | 5'-ATA**ATCGAT**CGCATGAAGGGCGCGCCAA-3' | For pMD19-T/KSK construction with Cla I site |
| P022 | pMD19-T/K-Spinach1-K (Mlu I) Rev | 5'-ATA**ACGCGT**CATGAGGCCCAGGTTGTTAATTAATTAT-3' | For pMD19-T/KSK construction with Mlu I site |
| P023 | pMD19-T/K-Spinach1-K (Eag I) For | 5'-ATA**CGGCCG**CGCATGAAGGGCGCGCCAA-3' | For pMD19-T/KSK construction with Eag Isite |
| P024 | pMD19-T/K-Spinach1-K (BspE I) Rev | 5'-ATA**TCCGGA**CATGAGGCCCAGGTTGTTAATTAATTAT-3' | For pMD19-T/KSK construction with BspE I site |
| P025 | pMD19-T/K-Spinach1-K (BspE I) For | 5'-ATA**TCCGGA**CGCATGAAGGGCGCGCCAA-3' | For pMD19-T/KSK construction with BspE I site |
| P026 | pMD19-T/K-Spinach1-K (EcoR V) Rev | 5'-ATA**GATATC**CATGAGGCCCAGGTTGTTAATTAATTAT-3' | For pMD19-T/KSK construct with EcoR V site |
| P027 | pMD19-T/K-Spinach1-K (EcoR V) For | 5'-ATA**GATATC**CGCATGAAGGGCGCGCCAA-3' | For pMD19-T/KSK construction with EcoR V site |
| P028 | pMD19-T/K-Spinach1-K (Sal I) Rev | 5'-ATA**GTCGAC**CATGAGGCCCAGGTTGTTAATTAATTAT-3' | For pMD19-T/KSK construction with Sal I site |
| P029 | OsU6:K-K For | 5'-ATGC**GATTGGGTCTTC**CCATCACCACCACGGAATCGATA-3' | For OsU6:KK construction with Bbs I site |
| P030 | OsU6:K-K Rev | 5'-TACG**AAACAGGTCTTC**CATCGGCGGTCGACGATATCAT-3' | For OsU6:KK construction with Bbs I site |
| P031 | OsU6:K-Spinach1-K For | 5'-ATGC**GATTGGGTCTTC**CGCATGAAGGGCGCGCCA-3' | For OsU6:KSK construction with Bbs I site |
| P032 | OsU6:K-Spinach1-K Rev | 5'-TACG**AAACAGGTCTTC**CATGAGGCCCAGGTTGTTAATTAAT-3' | For OsU6:KSK construction with Bbs I site |
| P033 | pEAQ-HT/K-Spinach1-K (K-K)-RT For | 5'-GGTTTTCGAACTTGGAGAAAG-3' | RT-PCR primers for pEAQ-HT/KSK (or KK) transgenic plants |
| P034 | pEAQ-HT/K-Spinach1-K (K-K)-RT Rev | 5'-ACACCGAATAACAGTAAATTCAAAC-3' | RT-PCR primers for pEAQ-HT/KSK (or KK) transgenic plants |
| P035 | pCVA sequence For | 5'-GGCGGCCGCTCTAGTGGAT-3' | For pCVA/KK (or KSK) sequencing |
| P036 | pCVA sequence Rev | 5'-CCCCTATATTTCAGGATACAACGC-3' | For pCVA/KK (or KSK) sequencing |
| P037 | pMD19-T M13-47 | 5'-CGCCAGGGTTTTCCCAGTCACGAC-3' | For pMD19-T/KK (or KSKs) sequencing |
| P038 | pMD19-T RV-M | 5'-GAGCGGATAACAATTTCACACAGG-3' | For pMD19-T/KK (or KSKs) sequencing |
| P039 | pEAQ-HT Sequence For | 5'-GGTGTGGTCTTGGGAAAAGA-3' | For pEAQ-HT/KK (or KSK) sequencing |
| P040 | pEAQ-HT Sequence Rev | 5'-CTGAAGGGACGACCTGCTAA-3' | For pEAQ-HT/KK (or KSK) sequencing |
| P041 | PVX Sequence For | 5'-CAGTGTTGGCTTGCAAACTAG-3' | For PVX/KK (or KSK) sequencing |
| P042 | PVX Sequence Rev | 5'-ACTATGGCACGGGCTGTACTAAAG-3' | For PVX/KK (or KSK) sequencing |
